# Behavioral effects of targeting the central thalamus and pulvinar with transcranial ultrasound stimulation in healthy volunteers

**DOI:** 10.1101/2025.02.27.640692

**Authors:** Amber R. Hopkins, Joshua A. Cain, Elizabeth A. White, Stephanie M. Abakians, Martin M. Monti

## Abstract

We present the first causal evidence in the healthy human brain linking the central thalamus to vigilance and the pulvinar to visuospatial attention using low-intensity transcranial focused ultrasound stimulation (tFUS). In a within-subjects, counterbalanced design, 27 healthy volunteers completed the Psychomotor Vigilance Task (PVT), a vigilance task, and the Egly-Driver Task (EDT), which assesses visuospatial attention, before and after central thalamus, pulvinar, and sham sonication with tFUS. Central thalamic sonication significantly impaired performance on both tasks in a manner consistent with decreases in arousal. Targeting the pulvinar with tFUS resulted in more subtle changes in visuospatial attention. Participants were less reactive to visual stimuli presented in the visual field contralateral to the affected pulvinar. These results demonstrate that tFUS can elicit measurable behavioral changes in healthy volunteers and underscores its potential as a high-resolution tool for non-invasive brain mapping, capable of differentiating functional contributions of thalamic regions only millimeters apart.

## Introduction

The thalamus and its interactions with the cortex play a central role in perception, attention, arousal, and consciousness. However, because common neurostimulation techniques that are safe for use with healthy volunteers, such as transcranial magnetic stimulation (TMS), cannot directly alter activity in deep brain structures like the thalamus, current knowledge about the contributions of different parts of the thalamus to healthy human cognition relies on correlational neuroimaging studies with healthy volunteers or lesion and invasive neurostimulation work with non-human animal models and human patient populations [1–5]. The emerging technique of low-intensity focused transcranial ultrasound stimulation (tFUS) addresses this gap, allowing for the safe [6–8] and non-invasive modulation of cortical or subcortical tissue with high spatial precision [9] in the healthy human brain [10]. In this work, we leverage tFUS to dissociate the roles of two functionally distinct parts of the thalamus in healthy volunteers, including the central thalamus and the pulvinar.

A growing body of work implicates the central thalamus, which includes the intralaminar thalamic nuclei and adjacent areas, in arousal regulation and consciousness [1, 11]. The central thalamus not only bridges the brainstem and cortex as part of the ascending reticular activating system [12–14] but also exhibits widespread and reciprocal connections with the cortex that put it in position to facilitate cortical synchrony and electrocortical arousal [1, 11, 15–18]. Indeed, invasive electrical stimulation of the central thalamus in awake non-human primates can increase arousal [19] or induce lapses in consciousness [20], depending on the parameters. Several studies also demonstrate that behavioral and electrocortical markers of arousal can be restored in anesthetized rodents [3, 4, 21] and non-human primates [1, 22–24] by delivering invasive electrical stimulation to the central thalamus, as well. Moreover, the application of invasive deep brain stimulation, and more recently tFUS, to the central thalamus in human patients with a disorder of consciousness (DOC) can bring some improvements in their behavioral responsiveness [25].

In contrast, the pulvinar, which subsumes the posterior third of the thalamus, shows interactions with the cortex that are consistent with a role in visual processing and visuospatial attention. The pulvinar is reciprocally connected with both the visual cortices and higher-order frontal and parietal areas [26–31]. Visual responses modulated by attention can be recorded from the pulvinar [32–34] with enhanced pulvinar activation observed under engaged visual attention [35, 36]. The pulvinar has been shown to synchronize cortical activity following attention, regulating the transmission of information between cortical areas according to attentional demands [37]. In addition, neural activity propagates from the pulvinar to cortex during states of engaged visual attention [38]. Moreover, lesions and deactivation of the pulvinar have been associated with deficits of visuospatial attention in humans [39–42] and non-human primates [2, 43–45], especially for the contralateral visual field and when stimuli in multiple locations compete for attention [2, 39, 42].

Leveraging the capability of tFUS to modulate subcortical tissue, we test causal roles of the central thalamus and the pulvinar in healthy human vigilance and visuospatial attention. In a within-subjects, sham-controlled, and counterbalanced design, 27 healthy volunteers completed the Psychomotor Vigilance Task (PVT), which assess vigilance [46], and the Egly-Driver Task (EDT), a visuospatial attention task [47], before and after central thalamic, pulvinar, and sham sonication with tFUS. Targeting the central thalamus with tFUS impaired performance on both tasks in a manner consistent with a reduction in arousal. Pulvinar sonication resulted in more subtle deficits in visuospatial attention. Specifically, participants were less responsive to visual stimuli presented in the visual field contralateral to the targeted pulvinar. These results constitute the first causal evidence in the healthy brain for the role of the central thalamus and pulvinar, underscoring the potential of tFUS as a high-resolution tool for causal brain mapping of deep brain regions in healthy volunteers.

## Results

### Psychomotor Vigilance Task (PVT)

Mixed-effects models testing the three-way interaction between the effects of ultrasound condition (sham, pulvinar, and central thalamic sonication), block (before and after sonication), and minutes into the task were used to examine whether targeting the central thalamus or pulvinar with tFUS altered behavioral markers of vigilance during the PVT [46], including response time, response speed, the slowest 10% of responses, and lapses in attention. Descriptive statistics for the data used in the models are presented in Fig. 2A, D, and G. Key results are presented in the main text. Model summaries and additional results are described in the Supplemental Material.

**Figure 1.**
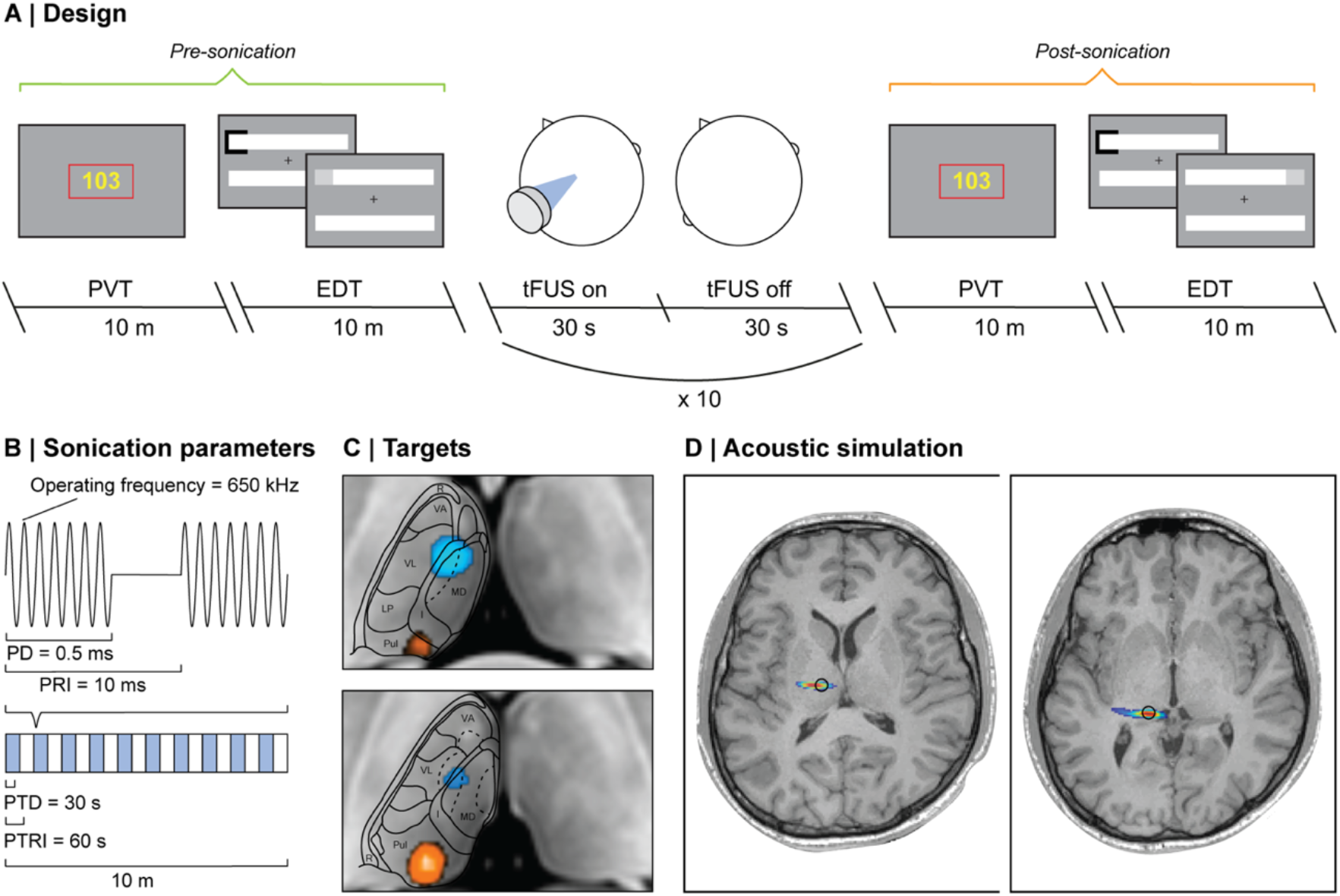
Methods (A): Session schematic. Participants complete the PVT and EDT before and after sham, pulvinar, and central thalamic sonication (in separate sessions and in a counterbalanced order across participants). During the PVT, participants maintained central fixation and pressed a button as quickly as possible when a visual cue (yellow millisecond counter) appeared. During the EDT, participants maintained central fixation and pressed a button if they detected a visual target. On main trials, a spatial cue (black square) indicated where the target was most likely to appear (75% cue validity). On catch trials a spatial cue appeared but no visual target followed. (B): Sonication regime. 10 blocks of 30 seconds of sonification (blue) and 30 seconds of rest (white), corresponding to a 30-second pulse train duration (PTD) and 60-second pulse train repetition interval (PTRI). We used a 0.5 ms pulse duration (PD) and 10 ms pulse repetition interval (PRI), resulting in a 5% (PD/PRI) duty cycle (DC) and a 100 Hz (1/PRI) pulse repetition frequency (PRF). The I_spta.3_ was ≤720 mW/cm^2^ and the I_sppa.3_ was ≤14.40 W/cm^2^. (C): Ultrasound targets. Heatmap showing tFUS aiming masks for the central thalamus (blue) and pulvinar (orange) across participants in standard space. 5-mm spheres were drawn in standard space in each target region and then warped onto the T1 image of each participant to create a tFUS aiming mask. (D): Simulated acoustic effects when targeting left central thalamus (left) and pulvinar (right) for an example participant. The focus of the ultrasound is colored by the amount of acoustic pressure achieved, from 50% of the maximum pressure (blue) to the maximum pressure (red). The left central thalamus (left) and pulvinar (right) are outlined in black according to the participant’s tFUS aiming mask.

**Figure 2.**
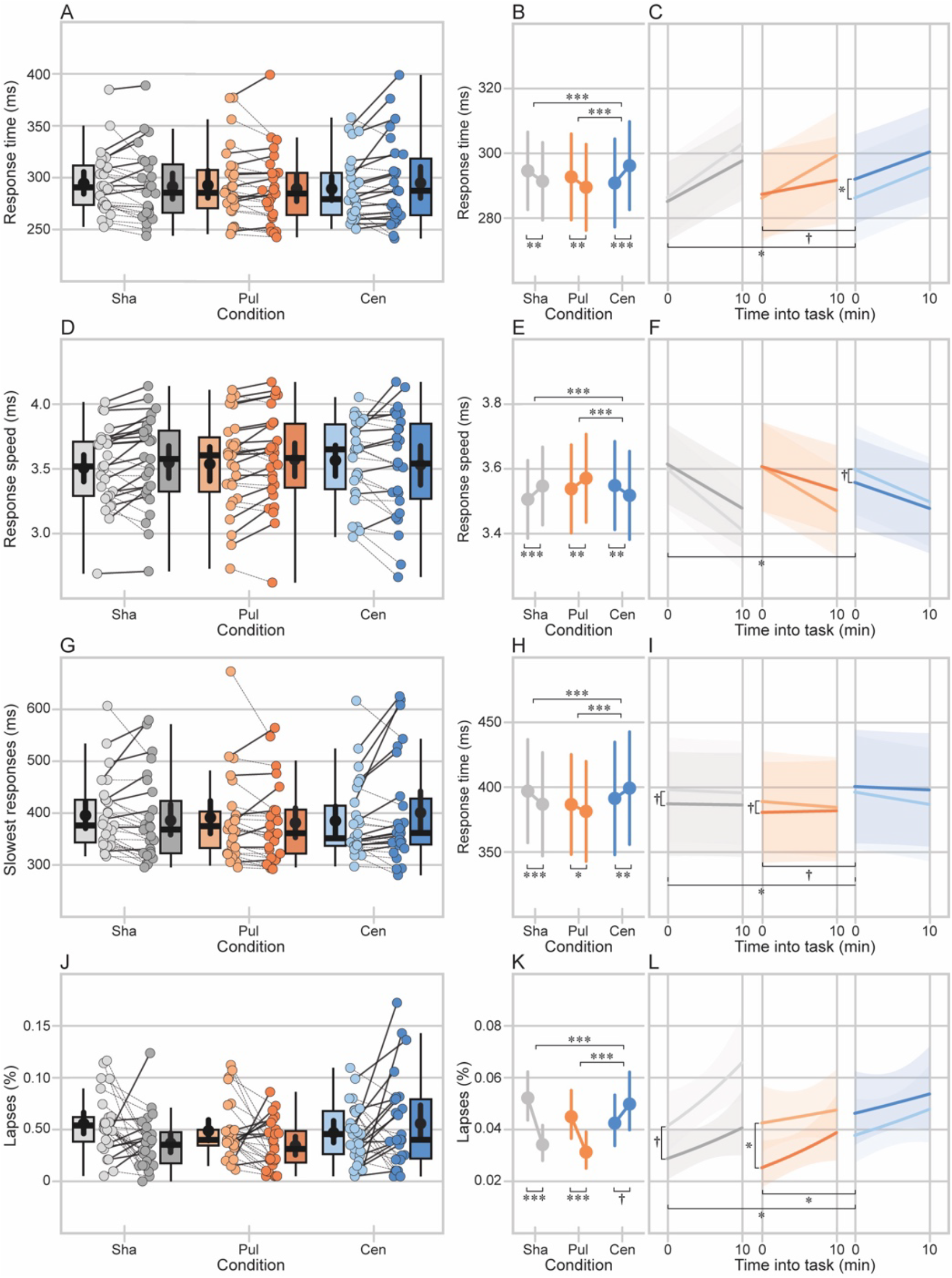
Central thalamic sonication reduced vigilance during the Psychomotor Vigilance Task (PVT). (A, D & G): Descriptive statistics for the data used in the model. Distribution of response times (A), response speeds (D), and lapses (G) before and after sonication (left to right) in the sham (left), pulvinar (middle) and central thalamus (right) conditions. Boxes show first quartile, median, and third quartile. Whiskers span the datapoints within 1.5 × the interquartile range (IQR) from the lower and upper hinges. Black points and bars show the means across participants and 95% confidence intervals, respectively. Individual points are participant means. Lines show each individual participant’s change in the outcome across blocks and are dashed or slide if that participant showed a higher value of the outcome before or after sonication, respectively. (B-C, E-F, & H-I): Mixed-effects modeling results. (B, E, & H): Estimated marginal means for response time (C), response speed (D), and predicted probability of lapsing (E) before and after sonication (left to right) for each ultrasound condition. Bars show 95% confidence intervals. (C, F, & I): Predicted response time (C), response speed (D), and probability of lapsing (E) before (light) and after (dark) for each ultrasound condition over the duration of the task (in minutes). Bands show 95% confidence intervals. Abbreviations: Sha, sham; Pul; pulvinar, Cen; central thalamus. † p < 0.10 before and/or after correction. * p < 0.05, ** p < 0.01, *** p < 0.001 after correction.

### Response time

Participants took longer to respond to the visual cue following central thalamic sonication compared to sham and pulvinar sonication (see Fig. 2B). This was evident at task onset and persisted for the duration of the task (see Fig. 2C). On average, response times decreased by 3.22 ms (SE = 0.98, 95% CI = [-5.13, −1.30], z = −3.29, p = 0.001, p_adj_ = 0.001) and 3.20 ms (SE = 0.98, 95% CI = [−5.12, −1.28], z = −3.27, p = 0.001, p_adj_ = 0.001) after sham and pulvinar sonication, respectively, but increased by 5.34 ms (SE = 1.00, 95% CI = [3.38, 7.30], z = 5.35, p < 0.001, p_adj_ < 0.001) after central thalamic sonication (see Fig. 2B). The change in average response times after sonication was 8.56 ms (SE = 1.40, 95% CI = [5.82, 11.30], z = 6.13, p < 0.001, p_adj_ < 0.001) and 8.55 ms (SE = 1.40, 95% CI = [5.80, 11.29], z = 6.11, p < 0.001, p_adj_ < 0.001) greater for the central thalamus condition compared to the sham and pulvinar conditions, respectively. Response times at task onset (0 minutes into the task) did not change after sham (−1.29 ms, SE = 1.93, 95% CI = [−5.07, 2.50], z = −0.67, p = 0.51, p_adj_ = 0.65) or pulvinar (1.11 ms, SE = 1.94, 95% CI = [−2.68, 4.91], z = 0.57, p = 0.57, p_adj_ = 0.65) sonication, but increased by 5.80 ms (SE = 1.98, 95% CI = [1.93, 9.68], z = 2.94, p = 0.003, p_adj_ = 0.024) after central thalamic sonication (see Fig. 2C). The change in response times at task onset after sonication was 7.09 ms (SE = 2.76, 95% CI = [1.67, 12.51], t = 2.57, p = 0.010) and 4.69 ms (SE = 2.77, 95% CI = [−0.73, 10.11], z = 1.70, p = 0.089, p_adj_ = 0.21) greater for the central thalamus condition compared to the sham and pulvinar conditions, respectively. Central thalamic sonication did not alter the average change in response time per minute in the task compared to sham (0.30 ms, SE = 0.49, 95% CI = [−0.65, 1.25], t = 0.62, p = 0.54) or pulvinar (0.78 ms, SE = 0.49, 95% CI = [−0.17, 1.74], z = 1.62, p = 0.10, p_adj_ = 0.21) sonication.

### Response speed

Responses slowed after central thalamic sonication compared to sham and pulvinar sonication (see Fig. 2E). This started at task onset and persisted for the duration of the task (see Fig. 2F). On average, response speeds increased by 0.04 ms (SE = 0.01, 95% CI = [0.02, 0.06], z = 4.21, p < 0.001, p_adj_ < 0.001) and 0.03 ms (SE = 0.01, 95% CI = [0.01, 0.05], z = 3.35, p = 0.001, p_adj_ = 0.002) after sham and pulvinar sonication, respectively, but decreased by 0.03 ms (SE = 0.01, 95% CI = [−0.05, −0.01], z = −3.04, p = 0.002, p_adj_ = 0.002) after central thalamic sonication (see Fig. 2E). The change in average response speeds after sonication was 0.07 ms (SE = 0.01, 95% CI = [−0.10, −0.04], z = −5.12, p < 0.001, p_adj_ < 0.001) and 0.06 ms (SE = 0.01, 95% CI = [−0.09, −0.04], z = −4.51, p < 0.001, p_adj_ < 0.001) lower for the central thalamus condition compared to the sham and pulvinar conditions, respectively. Response speed at task onset (0 minutes into the task) did not change after sham sonication (0.02 ms, SE = 0.02, 95% CI = [−0.02, 0.05], z = 0.80, p = 0.42, p_adj_ = 0.56) or pulvinar sonication (0.00 ms, SE = 0.02, 95% CI = [−0.04, 0.04], z = 0.07, p = 0.94, p_adj_ = 0.94) but decreased by 0.04 ms (SE = 0.02, 95% CI = [−0.08, 0.00], z = −2.02, p = 0.04, p_adj_ = 0.24) after central thalamic sonication (see Fig. 2F). The change in response speed at task onset after sonication was 0.06 ms (SE = 0.03, 95% CI = [−0.11, 0.00], t = −2.01, p = 0.045) lower for the central thalamus condition than sham. Central thalamic sonication did not alter the change in response speed per minute into the task compared to sham (0.00 ms, SE = 0.00, 95% CI = [−0.01, 0.01], t = −0.67, p = 0.50) and pulvinar (0.00 ms, SE = 0.49, 95% CI = [−0.01, 0.01], z = 0.91, p = 0.36, p_adj_ = 0.56) sonication.

### Slowest 10% of responses

The slowest 10% of responses increased after central thalamic sonication compared to sham and pulvinar sonication (see Fig. 2H). This started at task onset and persisted for the duration of the task (see Fig. 2I). On average, the slowest responses decreased by 10.11 ms (SE = 2.33, 95% CI = [−14.67, −5.55], z = −4.35, p < 0.001, p_adj_ < 0.001) and 5.32 ms (SE = 2.38, 95% CI = [−9.98, −0.65], z = −2.23, p = 0.025, p_adj_ = 0.030) after sham and pulvinar sonication, respectively, but increased by 7.96 ms (SE = 2.38, 95% CI = [3.19, 12.73], z = 3.27, p = 0.001, p_adj_ = 0.002) after central thalamic sonication (see Fig. 2H). The change in the average slowest responses after sonication was 18.07 ms (SE = 3.37, 95% CI = [11.47, 24.67], z = 5.37, p < 0.001, p_adj_ < 0.001) and 13.28 ms (SE = 3.40, 95% CI = [6.60, 19.95], z = 3.90, p < 0.001, p_adj_ < 0.001) greater for the central thalamus condition compared to the sham and pulvinar conditions, respectively. The slowest responses at task onset (0 minutes into the task) decreased by 10.95 ms (SE = 4.79, 95% CI = [−20.34, −1.57], z = −2.29, p = 0.022, p_adj_ = 0.17) and 8.31 ms (SE = 4.99, 95% CI = [−18.09, 1.46], z = −1.67, p = 0.095, p_adj_ = 0.25) after sham and pulvinar sonication, respectively, but not after central thalamic sonication (4.34 ms, SE = 5.07, 95% CI = [−1.28, 21.73], z = 0.86, p = 0.39, p_adj_ = 0.65; see Fig. 2I). The change in the slowest responses at task onset after sonication was 15.30 ms (SE = 6.97, 95% CI = [1.62, 28.97], t = 2.88, p = 0.028) and 12.66 ms (SE = 7.11, 95% CI = [−1.28, 26.59], z = 1.78, p = 0.075, p_adj_ = 0.25) greater for the central thalamus condition compared to the sham and pulvinar conditions, respectively (see Fig. 2I). Central thalamic sonication did not alter the average change in the slowest responses per minute into the task beyond sham (0.53 ms, SE = 1.18, 95% CI = [−1.78, 2.84], t = 0.45, p = 0.65) and pulvinar (0.12 ms, SE = 1.20, 95% CI = [−2.23, 2.46], z = 0.10, p = 0.92, p_adj_ = 0.92) sonication.

### Lapses in attention

Lapses in attention became more frequent after central thalamic sonication compared to sham and pulvinar sonication (see Fig. 2K). This started at task onset and persisted for the duration of the task (see Fig. 2L). On average, participants were 0.64 times as likely to lapse after sham sonication (95% CI = [0.53, 0.77], z = −4.66, p < 0.001, p_adj_ < 0.001) and 0.69 times as likely to lapse after pulvinar sonication (95% CI = [0.57, 0.83], z = −3.80, p < 0.001, p_adj_ < 0.001). However, they were 1.18 times more likely to lapse after central thalamic sonication (95% CI = [0.99, 1.41], z = 1.84, p = 0.066, p_adj_ = 0.079; see Fig. 2K). The odds of lapsing after sonication were 1.84 times (95% CI = [1.42, 2.38], z = 4.64, p < 0.001, p_adj_ < 0.001) and 1.72 times (95% CI = [1.32, 2.24], z = 4.04, p < 0.001, p_adj_ < 0.001) higher for the central thalamus condition compared to the sham and pulvinar conditions, respectively. At task onset (0 minutes into the task), participants were 0.69 times as likely to lapse after sham sonication (95% CI = [0.47, 1.01], z = −1.93, p = 0.054, p_adj_ = 0.14) and 0.59 times as likely after pulvinar sonication (95% CI = [0.39, 0.87], z = −2.65, p = 0.008, p_adj_ = 0.032). However, participants were 1.24 times more likely to lapse at task onset after central thalamic sonication (95% CI = [0.86, 1.77], z = 1.15, p = 0.25, p_adj_ = 0.50). The odds of lapsing at task onset after sonication were 1.80 times (95% CI = [1.06, 3.04], z = 2.19, p = 0.028) and 2.11 times (95% CI = [1.24, 3.60], z = 2.74, p = 0.006, p_adj_ = 0.032) times higher for the central thalamus condition compared to the sham and pulvinar conditions, respectively. Central thalamic sonication did not alter the change in the odds of lapsing per minute into the task beyond sham (odds ratio = 1.00, 95% CI = [0.92, 1.10], z = 0.10, p = 0.92) and pulvinar (odds ratio = 0.96, 95% CI = [0.88, 1.05], z = −0.89, p = 0.37, p_adj_ = 0.50) sonication.

### Egly-Driver Task (EDT)

Mixed-effects models were used to examine whether targeting the pulvinar or central thalamus with tFUS affected visuospatial attention during EDT [47], specifically the frequency of correct responses and response time. Descriptive statistics for the cleaned data used in the models are presented in the Supplemental Material. Additional results are provided in the Supplemental Material.

### Accuracy

Participants detected validly cued visual targets on main trials less often after central thalamic sonication, whether the spatial cue and visual target appeared ipsilaterally or contralaterally to the affected thalamus (see Fig. 3B). Specifically, participants were 0.69 times (95% CI = [0.56, 0.85], z = −3.45, p = 0.001, p_adj_ = 0.015) as likely to detect validly cued ipsilateral targets after central thalamic sonication compared to sham. Similarly, participants were 0.69 times (95% CI = [0.55, 0.87], z = −3.21, p = 0.001, p_adj_ = 0.015) and 0.79 times (95% CI = [0.63, 0.98], z = −2.17, p = 0.030, p_adj_ = 0.23) as likely to detect validly cued contralateral targets after central thalamic sonication compared to sham and pulvinar sonication, respectively.

**Figure 3.**
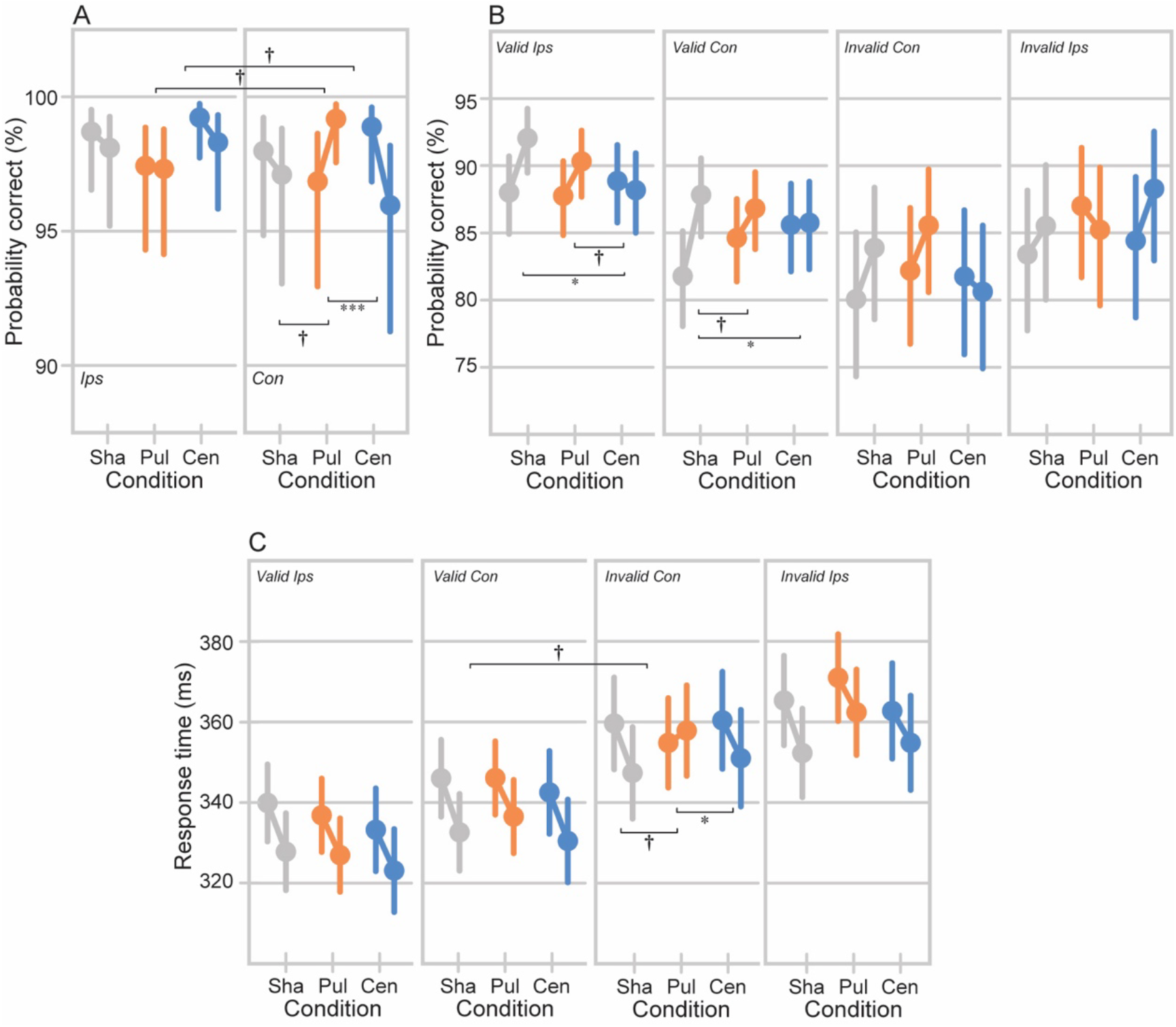
Mixed-effects modeling results for the Edgly-Driver Task (EDT). Bars show 95% confidence intervals. (A): Estimated marginal means for the predicted probability of responding correctly on catch trials (‘correct rejections’) with ipsilateral (left) and contralateral (right) spatial cues for each condition before and after sonication (left to right). (B & C): Estimated marginal means for the predicted probability of responding correctly (B) and response times (C) on main trials for each combination of cue validity and visual field the target appeared in, including validly cued ipsilateral targets, validly cued contralateral targets, invalidly cued ipsilateral targets, and invalidly cued contralateral targets. Abbreviations: Sha, sham; Pul, pulvinar; Cen, central thalamus; Ips, ipsilateral to the targeted regions; Con, contralateral to the targeted regions. † p < 0.10 before and/or after correction. * p < 0.05, ** p < 0.01, *** p < 0.001 after correction.

Participants were less reactive to contralateral spatial cues on catch trials after pulvinar sonication compared to sham and central thalamic sonication (see Fig. 3A), as indicated by more correct responses (‘correct rejections’) from fewer button presses. Participants were 2.76 times (95% CI = [0.97, 7.87], z = 1.90, p = 0.058, p_adj_ = 0.17) more likely to correctly reject on catch trials with contralateral spatial cues after pulvinar sonication compared to sham sonication, and 0.20 times (95% CI = [0.06, 0.62], z = −2.78, p = 0.005, p_adj_ = 0.045) as likely after central thalamic sonication compared to pulvinar sonication. Additionally, participants were 2.51 times (95% CI = [0.90, 6.95], z = 1.77, p = 0.077, p_adj_ = 0.17) more likely to correctly reject on catch trials with contralateral spatial cues than ipsilateral ones after pulvinar sonication but not after sham sonication (odds ratio = 1.09, 95% CI = [0.39, 3.05], z = 0.17, p = 0.88) or central thalamic sonication (odds ratio = 0.55, 95% CI = [0.18, 1.72], z = −1.02, p = 0.31, p_adj_ = 0.46). Participants were 2.29 times (95% CI = [0.54, 9.72], z = 1.12, p = 0.26) more likely to correctly reject catch trials with contralateral spatial cues than ipsilateral ones after pulvinar sonication compared to sham sonication, and 0.22 times (95% CI = [0.05, 0.92], z = −1.94, p = 0.05, p_adj_ = 0.19) as likely after central thalamic sonication compared to pulvinar sonication.

Pulvinar sonication also interfered with contralateral visual target detection during main trials (see Fig. 3B). Participants were 0.79 times (95% CI = [0.64, 0.98], z = −2.19, p = 0.029, p_adj_ = 0.23) as likely to detect validly cued ipsilateral targets after pulvinar sonication compared to sham sonication. In addition, participants were 0.78 times (95% CI = [0.46, 1.32], z = −0.92, p = 0.36) as likely to detect invalidly cued contralateral targets after pulvinar sonication compared to sham, and 1.45 times (95% CI = [0.83, 2.53], z = 1.31, p = 0.19, p_adj_ = 0.43) more likely compared to central thalamic sonication. Finally, while participants were 1.38 times (95% CI = [0.82, 2.32], z = 1.20, p = 0.23, p_adj_ = 0.43) more likely to detect invalidly cued ipsilateral targets than invalidly cued contralateral targets after pulvinar sonication, this effect did not reliably differ from sham (1.26, 95% CI = [0.63, 2.53], z = 0.66, p = 0.51) or central thalamic, (0.53, 95% CI = [0.26, 1.09], z = −1.72, p = 0.086, p_adj_ = 0.36) sonication. While many comparisons involving pulvinar sonication were marginally significant after correction or did not pass correction, the confidence intervals suggest that differences could become more pronounced with additional data.

### Response time

Participants took longer to respond to visual targets when the cue direct their attention ipsilaterally, but the target appeared contralaterally (see Fig. 3C). Participants took 15.30 ms (SE = 6.96, 95% CI = [1.66, 28.95], t = 2.20, p = 0.028) and 12.43 ms (SE = 6.96, 95% CI = [−1.21, 26.06], z = 1.40, p = 0.074, p_adj_ = 0.55) longer to respond to invalidly cued contralateral visual targets after pulvinar sonication compared to sham and central thalamic sonication, respectively. Participants took 12.62 ms (SE = 5.33, 95% CI = [2.48, 23.36], z = 2.43, p = 0.015, p_adj_ = 0. 26) longer to respond to invalidly cued contralateral targets than validly cued contralateral targets after pulvinar sonication. This was 13.08 ms (SE = 7.43, 95% CI = [−1.47, 27.64], t = 1.76, p = 0.078) and 12.20 ms (SE = 7.42, 95% CI = [−2.34, 26.73], z = 1.64, p = 0.10, p_adj_ = 0.55) greater compared to sham and central thalamic sonication, respectively. Similarly, participants took 12.62 ms (SE = 5.31, 95% CI = [−2.21, 23.02], z = 2.38, p = 0.017, p_adj_ = 0.26) longer to respond to invalidly cued contralateral targets than validly cued ipsilateral targets after pulvinar sonication. This was 11.47 ms (SE = 7.40, 95% CI = [− 3.04, 25.97], t = 1.55, p = 0.12) and 9.94 ms (SE = 7.39, 95% CI = [−4.55, 24.43], z = 1.34, p = 0.18, p_adj_ = 0.60) greater compared to sham and central thalamic sonication, respectively. The change in response time after pulvinar sonication was 10.48 ms (SE = 6.66, 95% CI = [2.57, 23.53], t = 1.57, p = 0.12) greater for invalidly cued contralateral visual targets than invalidly cued ipsilateral visual targets. And this was 10.85 ms (SE = 9.36, 95% CI = [−7.48, 29.19], t = 1.16, p = 0.25) and 13.09 ms (SE = 9.40, 95% CI = [− 5.33, 31.52], z = 1.39, p = 0.16, p_adj_ = 0.60) greater than what was observed after sham and central thalamic sonication, respectively. While many comparisons were not significant or did not pass correction, the direction and range of the confidence intervals indicate these differences may become more pronounced with additional data.

## Discussion

Targeting the central thalamus with tFUS slowed responses and increased lapses during the PVT as well as reduced visual target detection rates during the EDT, which is consistent with a reduction in arousal level. Pulvinar sonication resulted in more specific deficits in visuospatial attention. Participants were less reactive to spatial cues on catch trials when they appeared in the visual field contralateral to the affected pulvinar during the EDT. In addition, the participants failed to detect, and took longer to respond to, contralateral visual targets on main trials when the spatial cue had directed their attention ipsilaterally during the EDT. These results provide the first causal evidence in the healthy human brain that the central thalamus and pulvinar are involved in arousal and visuospatial attention, respectively. However, this work is also the first to demonstrate that tFUS can modulate behavior over a period of at least 20 minutes and is spatially precise enough to elicit different behavioral effects when applied to subcortical regions that are only millimeters apart in the same individuals.

Reduced vigilance under diminished states of arousal like sleep deprivation [48] is reliably accompanied by slower responses [49, 50] and more frequent lapses [49, 51] during the PVT. We achieved similar results by targeting the central thalamus with tFUS in this work. Participants responded more slowly and showed lapses in attention more often during the PVT after central thalamic sonication compared to sham and pulvinar sonication (see Fig. 2). These behavioral changes could be consistent with a suppression of the central thalamus. In sleep deprived healthy volunteers, lapses are associated with reduced thalamic activation while non-lapse periods are accompanied by elevated thalamic activation [52]. Indeed, a recent study from our group targeted anterior parts of the left thalamus in awake healthy volunteers with tFUS at the same parameters we used in this work and demonstrates decreases in blood-oxygen level dependent (BOLD) signals during sonication and blood perfusion after sonication in the targeted thalamic regions but also in connected cortical areas in both hemispheres [53].

Thalamic activity during vigilance tasks in sleep deprived individuals reflects a complex interplay between the effects of sleep loss, which dampens arousal, and engagement in the vigilance, which recruits alertness [54]. At rest, thalamic activity is suppressed under sleep deprivation compared to rested wakefulness [55–58]. However, during vigilance tasks like the PVT, the thalamus shows greater activation under sleep deprivation than rested wakefulness, which indicates that the thalamus compensates for the dampening effects of sleep loss to facilitate engagement in the task [54, 56, 59]. Some work reports that these effects are stronger in more central parts of the thalamus, as well [54, 55]. If targeting the central thalamus with tFUS in our experiment emulates the effects of sleep deprivation, we might thus expect thalamic activity to be more pronounced during the PVT after central thalamic sonication. Future work could implement our experimental design in the MR environment to examine whether targeting the central thalamus with tFUS disrupts or amplifies the interplay of thalamic activity observed with vigilance tasks during sleep deprivation.

Where the alterations of behavioral markers of vigilance that we observed after targeting the central thalamus with tFUS differs from those reported for sleep deprivation involves the time-on-task effect (also called the vigilance decrement), whereby performance on a vigilance task worsens over the duration of the task due to factors like fatigue or boredom [60–63]. For the PVT, the time-on-task effect presents as a slowing of responses, an increase in lapses, and an increase in response time variability over the duration of the task. This can be observed during rested wakefulness but is stronger under states of diminished arousal like sleep deprivation [49, 62–65]. Our participants show the canonical time-on-task effect for the PVT. Their responses slowed and they showed lapses in attention more frequently over time into the task (see Fig. 2). However, the time-on-task effect was not altered by sonication in any of our ultrasound conditions (see the Supplemental Material). This may indicate central thalamic sonication in this work reduced arousal more generally but did not modulate the capacity to maintain vigilance over uninterrupted periods. There is some evidence that maintaining vigilance over uninterrupted periods recruits a predominantly right-lateralized network of cortical and subcortical structures, including right fronto-parietal areas and the right thalamus [66, 67]. Activity in these regions increases over longer durations of uninterrupted attentional demands [66] and patients with damage to them show stronger deficits in sustained attention than patients with lesions in their left hemisphere [68–72]. This could explain why we do not observe any changes in the time-on-task effect during the PVT after central thalamic sonication in our experiment. If maintaining vigilance over the task recruits a predominantly right-lateralized network, then we might not expect targeting the left central thalamus like we did in our experiment to alter how participants maintain vigilance over the duration of the task.

Pulvinar lesions and deactivation are associated with deficits in visuospatial attention in humans [39–42] and non-human primates [2, 43–45], especially toward stimuli in the visual field contralateral to the affected pulvinar. Behavioral impairments in human patients with pulvinar damage are especially strong when distractor stimuli compete for attention [39, 42]. For instance, one study used muscimol to inactivate the pulvinar in behaving non-human primates and observed diminished exploration of the contralesional visual field, especially when stimuli appeared in both visual fields [2]. We observed similar changes in visuospatial attention after targeting the pulvinar with tFUS. Participants were less reactive to contralateral spatial cues on catch trials during the EDT after pulvinar sonication, which is reflected by an increase in correct responses (‘correct rejections’) from fewer responses. In addition, participants took longer to respond to contralateral visual targets when a spatial cue had directed their attention to the ipsilateral visual field prior, competing for their attentional resources. While our results are consistent with the literature, they are statistically weak because the low number of catch trials and invalidly cued main trials in the abbreviated EDT we used degraded the statistical power for these comparisons. Future work administering the entire version of the EDT, before and after targeting the pulvinar with tFUS, will be necessary to make stronger conclusions about the effects of pulvinar sonication on visuospatial performance during the EDT.

By selecting cue-to-target intervals from a continuous range of values between 300 and 1100 ms during the EDT, fluctuations in spatial and object attention as a function of the interval between a spatial cue and visual target have been mapped to demonstrate that visuospatial attention is discontinuous, sampling the environment in theta-rhythmic cycles (3–8 Hz) [38, 73–75]. These cycles are thought to organize neural activity into alternating attentional states of engagement (heightened sensitivity to stimuli in the attended location) and disengagement (decreased sensitivity to stimuli in the attended location and increased probability of shifting attention to another location) [74]. While these theta-rhythmic cycles in visuospatial attention have been linked to theta oscillations in frontal and parietal cortices [74, 76], the pulvinar seems to play an active and coordinating role in them. A recent study assessed pulvino-cortical interactions during theta-rhythmic sampling by recording the frontal eye fields, lateral intraparietal area, and pulvinar in awake non-human primates completing the EDT [38]. The authors report that activity propagated from the pulvinar to the cortex during engaged states of attention and from the cortex to the pulvinar during disengaged states. We might thus expect suppressing the pulvinar with tFUS in this work to disrupt theta-rhythmic sampling of visuospatial attention during the EDT. However, because we used an abbreviated version of the EDT that used only (300 ms) and long (1100 ms) intervals between the spatial cue and visual target to ensure that task duration was held constant between our tasks, we cannot examine whether pulvinar sonication in our experiment altered the theta-rhythmic cycles of attention. Future work could administer the entire EDT, selecting cue-to-target intervals from the continuous range of values between 300 and 1100 ms, to test whether targeting the pulvinar with tFUS can disrupt the theta-rhythmic sampling of visuospatial attention.

These results join a rapidly growing body of work applying tFUS in the healthy human brain. A variety of online (during sonication) behavioral effects have been reported for tFUS so far. Reduced reaction times have been reported for simple response time tasks [77] and visuomotor tasks [78] while delivering ultrasound to the primary motor cortex. Applying tFUS to the primary somatosensory cortex can induce tactile sensations [79, 80] and improve sensory discrimination of tactile stimuli [81]. In another study, delivering ultrasound to unilateral sensory thalamus worsened performance on a tactile discrimination task [82]. tFUS of the inferior frontal gyrus has been reported to improve response inhibition [83]. Fewer errors in an anti-saccade task were reported while delivering tFUS to the dorsolateral prefrontal cortex in one study [84]. tFUS to the visual cortex has been shown to induce phosphenes [85]. Fewer studies report the offline (after sonication) behavioral effects of tFUS, especially for long durations. Alterations of mood have been reported for up to 30 min after applying ultrasound to the frontal cortex [86, 87], and another study reports altered pain thresholds for up to 10 minutes after anterior thalamic sonication [88]. The bulk of previous work reports the online (during sonication) behavioral effects from delivering tFUS to one cortical or subcortical region, and only one previous study reports offline behavioral changes after thalamic ultrasound (for up to 10 minutes) [88].

We demonstrate that tFUS can produce distinct behavioral effects by targeting different regions of the thalamus in the same healthy individuals that persist for at least 20 minutes without signs of dissipation. Specifically, stimulation of the central thalamus led to reduced behavioral markers of arousal and vigilance, while targeting the pulvinar resulted in selective impairments in visuospatial attention despite these regions being only millimeters apart. This highlights the high spatial specificity of tFUS and its capacity to modulate neurophysiological and behavioral functions. Each behavioral task lasted 10 minutes, and task order was counterbalanced across participants. There was minimal variation attributable to task order (see Supplemental Material), and no evidence of waning effects during the PVT task following central thalamic sonication. These observations indicate that the effects of our sonication protocol endured for the duration of the testing period. Our results demonstrate that tFUS can map and modulate the healthy human brain with exceptional spatial precision. Its ability to non-invasively and selectively influence subcortical structures opens new avenues for causal investigations of subcortical networks and the treatment of neurological disorders.

## Methods

All data analysis scripts, pre-processing pipelines, and anonymized raw data sets will be uploaded to an Open Science Framework (OSF) repository associated with this article.

### Participants

Right-handed and English-speaking adults (18 years of age or older) with normal or corrected-to-normal vision were recruited following procedures approved by the Institutional Review Board (IRB) at the University of California, Los Angeles (UCLA). Individuals were excluded from the experiment if they self-disclosed current or recent use of any psychoactive medications or recreational drugs (e.g., antidepressants or stimulants), a counter-indication for entering the magnetic resonance imaging (MRI) environment (e.g., MRI-incompatible implants), possible or planned pregnancy (in the short-term), or had hair longer than a ¼ of an inch over their left temporal region and were unwilling to shave it. Participants provided written informed consent and were compensated at a rate of $40 per hour for their participation. If participants voluntarily withdrew before completing the full protocol (loss to follow-up), then a new participant was recruited to replace them. Ten participants voluntarily withdrew from the study, including 3 after the MRI visit only and 7 after at least one sonication session. Twentyseven participants completed the entire protocol and are included in the analyses (age = 23.07 ± 3.33, 2 females).

### Design

A 3×2 within-subjects design was used to test whether targeting the pulvinar or central thalamus with tFUS affects behavioral markers of vigilance during the Psychomotor Vigilance Task (PVT) [46] or visuospatial attention during the Egly-Driver Task (EDT) [47]. Participants underwent an MRI scan followed by three sonication sessions, including one per ultrasound condition: sham, pulvinar, and central thalamus sonication. Session order was counterbalanced across participants, and participants were blinded to the ultrasound condition assigned to each session. The PVT and EDT were administered before and after sonication in a counterbalanced order across participants during every session. Sonication sessions were held at least 24 hours apart but around the same time of day for each participant.

### Magnetic resonance imaging (MRI)

MR data were acquired with a 3T Siemens Prisma fit MRI scanner at the UCLA Staglin IMHRO Center for Cognitive Neuroscience. For every participant, we acquired an anatomical image (T1-weighted) image for use with a frameless stereotactic neuronavigational system using a T1-weighted MPRAGE sequence with the following parameters: TR = 2,400 ms, TI = 1,060 ms, TE = 2.12 ms, flip angle = 88, voxel size = 1×1×1 mm isotropic, matrix size = 256×256, 192 slices. The raw T1-weighted data was converted from DICOM to NIFTI format using *dcm2niix* [89].

### tFUS target mask

We targeted the central thalamus with tFUS because of its association with arousal [1] and the pulvinar because of its role in visuospatial attention [27]. We targeted structures in the left hemisphere for all participants. Masks for each target region were generated in standard Montreal Neurological Institute (MNI) space by drawing 5-mm spheres over the pre-determined target voxels (see Fig. 1C). A tFUS target mask was created for each participant by warping each target region mask from standard structural (MNI) space onto their T1-weighted image using *applywarp* in the FMRIB Software Library (FSL; https://fsl.fmrib.ox.ac.uk; [90]). Linear and non-linear registrations between native structural (T1) and standard structural (MNI) images were performed using *flirt* and *fnirt* in FSL [91]. All steps were visually inspected and adjusted as needed to ensure anatomical plausibility. The tFUS target masks were overlayed on the individual T1 in a neuronavigational system during ultrasound transducer placement and aiming.

### Transcranial focused ultrasound stimulation (tFUS)

Low-intensity tFUS was administered using a BXPulsar 1002 drive system and an accompanying fixed-focal-length, single-element, and air-backed spherical transducer with a 80-mm fixed focal length, 61-mm aperture, and 650-kHz operating frequency (Brainsonix, Corp., USA) [92]. A Brainsight neuronavigation system (Rogue Research, Inc., Canada) was retrofitted for concurrent use with tFUS. Optical trackers placed on the participant’s head and a custom ultrasound transducer holder were co-registered with the participant’s T1 image in the Brainsight system, allowing us to visualize the transducer’s position relative to the participant’s brain in real time. Sham sonication was achieved by placing a pigmented gel pad that absorbs sound (Brainsonix Corp., USA) between the surface of the transducer and the head, preventing ultrasound transmission [93]. Acoustic coupling gel pads (Brainsonix Corp., USA) that promote ultrasound transmission to the head [92] were used during the genuine (pulvinar and central thalamus) sonication sessions, keeping set up was identical between sham and genuine sonication sessions. During sham sonication, the transducer was aimed at either the pulvinar or the central thalamus in a counterbalanced order across participants.

### Sonication regime

Ultrasound was delivered in an on/off block design including 10 blocks with 30 seconds of sonication and 30 seconds of rest. We used a 0.5 ms pulse duration (PD), 60 s pulse repetition interval (PRI), 5% duty cycle (DC), 100 Hz pulse repetition frequency (PRF), < 720 mW/cm^2^ I_spta.3_, and < 14.40 W/cm^2^ I_sppa.3_ based on previous work [53]. These intensities comply with the United States Food and Drug Administration (FDA) guidelines for diagnostic ultrasound [94] and have no known adverse thermal bioeffects [95, 96]. This regime was selected because previous work demonstrates that it has suppressive effects when applied to the thalamus [53].

### Procedure

The T1 image and tFUS aiming mask for the participant opened in the Brainsight neuronavigational system and a marker was generated in the center of the target region assigned to the session. The participant’s head was secured in a comfortable position using chin and head rests and they were instructed to remain as still as possible to minimize movement during transducer positioning and sonication. The transducer was first placed over the left temple (approximately 1/3 of the distance from the corner of the left eye to the left tragus and superior by 2 cm), which is the thinnest part of the temporal bone and ideal cranial entry-point for minimizing ultrasound disruption by the skull. Deviations in transducer position were made iteratively until the projected trajectory of the ultrasound beam passed through the target marker. The transducer was kept as flat against the side of the head as possible to deliver ultrasound perpendicular to the skull surface and minimize the scattering, reflection, or refraction of the ultrasound [97]. Once an adequate transducer position was achieved, aqueous ultrasound gel (Aquasonic Clear Ultrasound Transmission Gel; Parker Laboratories, Inc., USA) was applied to the region subsuming the diameter of the transducer such that no hair permeated the gel layer and air bubbles were pressed out [98, 99]. A thin layer of gel was also applied between the surface of the transducer and the gel pad with air bubbles smoothed out. If reaching the target region required tilting the transducer, then an angled gel pad that filled the expanding gap between the transducer and the scalp was used. After placing the transducer, additional gel was applied to fill any remaining concavities between the transducer and the scalp. Transducer position was verified continuously throughout sonication and adjustments were made as needed to ensure that the projected trajectory of the ultrasound passed through the target.

### Apparatus and stimuli

Stimuli and tasks were created and administered using PsychoPy (Version 2021.2.3; [100]). Participants were seated in upright position 150 cm away from the monitor with a button box in their right hand. Earplugs were provided to reduce auditory distractions.

### Psychomotor vigilance task (PVT)

The PVT assesses the capacity to maintain vigilance (also called sustained attention) over uninterrupted periods [46]. Participants maintained central fixation and pressed a button as quickly as possible when they saw a visual cue for 10 minutes without breaks. The visual cue was a yellow millisecond counter that appeared inside of a red rectangle at the center of the monitor at random 2 to 10 second inter-trial intervals. Response time was recorded every trial. The counter paused when the participant pressed the button but remained on the screen for one second before the trial ended, displaying the response time recorded on that trial. If the participant pressed the button before the visual cue was presented or too quickly afterward (response time ≤100 ms), then a false start message appeared. If the participant did not press the button within 30 seconds of the visual cue appearing, then the trial timed out and a beep played. Many outcomes can be derived from response times from the PVT. We selected mean response time, response speed (1/response time), the slowest 10% of response times, and unusually slow responses considered lapses in attention because of their sensitivity to states of diminished arousal like sleep deprivation [49, 101]. A time-on-task effect (also called a vigilance decrement) can be observed with the PVT whereby responses slow and lapses become more frequent over the duration of the task. Reduced vigilance arising from, for example, sleep deprivation, is associated with slower responses and more frequent lapses during the PVT overall but also exacerbates the time-on-task effects [50, 102–107].

### Egly-Driver Task (EDT)

The EDT assesses visuospatial attention [47]. Participants maintained central fixation and pressed a button as quickly as possible when they saw a visual target for 10 minutes. A 30-second break was provided every 2 minutes. Trial onset was marked by the presentation of two horizontal white bars, including one above fixation and another below it. After a variable delay between 400 and 800 ms, a spatial cue (black square) appeared at the end of one of the horizontal bars to direct attention to that location for 100 ms. On main trials (90% of the trials), a visual target (subtle change in contrast) appeared at the end of one of the horizontal bars for 100 ms after either a short (300-350 ms) or long (1000-1050 ms) interval. The visual target could appear in the cued location or another location. However, the spatial cue indicated the location where the visual target was most likely to appear (with 75% cue validity). The trial ended after a variable response window between 150 and 700 ms from target offset. On catch trials (10% of the trials), no visual target was presented, and the trial ended between 450 and 1750 ms after the spatial cue to match trial duration to the main trials.

Participants responded correctly or incorrectly on every trial. The correct response on main trials was press the button (‘hit’ and otherwise it was a ‘miss’). The correct response catch trials was not to press the button (‘correct rejection’ and otherwise it was a ‘false alarm’). Response time was recorded whenever the participant pressed the button. If the participant pressed the button on a main trial before the visual target appeared or too soon afterward (≤100 ms), then the trial was considered a false start and treated as incorrect response. The contrast of the visual target relative to the horizontal white bars was adjusted for each participant using a staircase procedure administered during their first sonication session. The staircase was identical to the main task described above, except that that the contrast of the visual target changed on every trial. The staircase followed a two-down, one-up rule whereby successful trials were followed by more difficult ones, decreasing the contrast of the visual target relative to the white bars until the participant failed to detect it, and then adjusting accordingly. The staircase ended when we identified the minimum contrast with which the participant correctly detected the visual target on 50% of the trials it appeared in. The final contrast value was saved and used in all EDT blocks for that participant.

Spatial attention is associated with enhanced sensory processing and behavioral responses for stimuli that appear in the attended region [108]. During the EDT, participants detect the visual target more often (and respond to it more quickly) on trials when the spatial cue correctly indicated the location it would appear in compared to trials when the spatial cue was invalid, demonstrating the behavioral benefit of visuospatial attention as well as the cost of needed to shift attention to another location [47]. By varying cue validity and the hemifield that the spatial cue and visual target appear in, we can also examine hemispheric differences in the behavioral benefits and costs associated with visuospatial attention with the EDT. Previous versions of the EDT used cue-to-target intervals that are randomly selected from a continuous range of values between 300 and 1100 ms [38, 73]. However, we administered an abbreviated version of the EDT that presents either short (300-350 ms) or long (1000-1050) intervals between the cue and target to avoid differences in task duration between the PVT and EDT that could limit the results, since we counterbalance order across the participants. We should still be able to determine whether sonication in any of the ultrasound conditions affects accuracy or response time, and whether there are hemispheric effects of sonication on visuospatial attention by varying cue and target location.

### Quantification and statistical analyses

Mixed-effects modeling was used to examine whether sonication in any of the ultrasound conditions affected behavioral markers of vigilance during the PVT and visuospatial attention during the EDT. We used mixed-effects models because of the repeated-measures structure of our data but also because they allow for the specification of random effects. Mixed-effects modeling was performed using the lme4 (Version 1.1.36; [109]), lmerTest (Version 3.1.3; [110]), and emmeans (Version 1.11.0; [111]), and DHARMa (Version 0.4.7; [112]) packages in R (Version 4.4.3; R Core Team [113]). Data cleaning and visualization was completed in Python (Version 3.11.11).

Random effects for participant and task order were included in all the mixed-effects models to better isolate the effects of sonication. Since participants underwent the ultrasound conditions in separate sessions on different days, we included a random intercept for participant with a random slope for condition (1 + condition | participant). The random slope for condition captures how the effect of ultrasound condition varies for each participant when all other fixed effects are at their reference levels (including block at pre-sonication), and thus accounts for individual differences in task performance before sonication between the sessions that could interfere with estimates of the fixed effects in the models. The random intercept for participant also accounts for having multiple observations per participant, since we are using unaggregated data from a repeated-measures design in the models. A random intercept for task order (1 | task order) was included to capture any differences in task performance that can be attributed to the order in which participants completed the behavioral tasks. The PVT and EDT were administered before and after sonication in every session but in a counterbalanced order across participants, separating the participants into two groups based on task order (PVT then EDT and EDT then PVT). Adding the random intercept for task order allows each group to have their own baseline values of the outcome variables in the models.

Mixed-effects logistic regressions were used for the binary outcomes, specifically lapses during the PVT and correct responses during the EDT. The response time on a PVT trial was either considered a lapse in attention or not, making lapses a binary variable in the unaggregated PVT data. Participants were either correct or not on an EDT trial, making correct responses a binary variable in the unaggregated EDT data. The fixed effects and estimated marginal means contrast outputs from these mixed-effects logistic regressions are reported in odds ratios, which indicate how much more or less likely an event is to occur in one group compared to another. An odds ratio of 1 indicates an equal likelihood of the event occurring in the groups. An odds ratio less than or greater than 1 indicates less or more likelihood of the event occurring in the first group than the second, respectively. Linear mixed-effects models were used for all continuous outcomes, including response time, the slowest 10% of responses, and response speed from the PVT as well as response time from the EDT. The fixed effects and estimated marginal means contrast outputs for the linear mixed-effects models are reported in the units of the dependent (outcome) variable in the model.

### Psychomotor vigilance task (PVT)

Mixed-effects modeling was used to examine whether sonication in any of the ultrasound conditions affected behavioral markers of vigilance during the PVT, including response time, response speed, the slowest 10% of response times, and lapses in attention for their sensitivity to states of diminished arousal like sleep deprivation [49]. Response speed was derived by dividing the response time recorded on every trial by 1000 and reciprocally transforming it. For the slowest 10% of responses, we restricted the data for each block to response times in the slowest 10% for that block. Lapses refer to lapses in attention and are traditionally defined as response times greater than 500 ms or twice the mean response time for an individual participant [4]. However, our participants never or rarely showed response times greater than 500 ms or twice their mean response time, even before trial cleaning (see the Supplemental Material). Instead, we combined the PVT data across all blocks for each participant and defined lapses as response times greater than the mean response time plus two standard deviations for that participant. Out of the 1129.37 trials (SD = 32.96, min = 1075, max = 1188) each participant completed in total on average, 55.15 were considered lapses in attention (SD = 7.98, min = 40, max = 69).

### Data cleaning

Individual trials were discarded if the response time recorded on that trial was a false start (≤100 ms) or a definite outlier relative to the other trials for that block and participant based on Tukey’s method (less than the first quartile minus 3 × the interquartile range (IQR) or greater than the third quartile plus 3 × the IQR) [114, 115]. This ensures that no erroneous response times within a block can drive the results. Participants completed 210.59 trials (SD = 5.29, min = 197, max = 226) per block, on average. 6.92 were false starts (SD = 4.72, min = 1, max = 25) and 6.06 were discarded as outliers (SD = 3.55, min = 1, max = 17). We were left with 197.42 trials per block for each participant, on average (SD = 7.33, min = 176, max = 214). To assess the influence of the participants on model estimates, we computed Cook’s distance for each participant for each mixed-effects model. Participants with Cook’s distance values substantially greater than the others (>4 × the mean) were considered influential and their data was examined further. Whole sessions from a participant were left out of the mixed-effects models if they heavily influenced the model estimates and their data was unusual. One session from one participant was left out of the models for response time, response speed, and lapses in attention. Three sessions from three separate participants were left out of the model for the slowest 10% of responses. All other data from these participants were used in the mixed-effects models to retain as much data as possible but also to improve estimates of the random effects in the models. See the Supplemental Material for additional information on participant exclusion.

### Analysis

Separate mixed-effects models were used to examine whether sonication in any of the ultrasound conditions affected response time, response speed, the slowest 10% of responses, or lapses during the PVT. Linear mixed-effects models were used for response time, response speed, and the slowest 10% of response times because they are continuous variables in the unaggregated data. A mixed-effects logistic regression was used for lapses because an individual trial was either considered a lapse in attention or not, making lapses a binary variable in the unaggregated data. All models shared the following specification: outcome ~ condition × block × minute + (1 + condition | participant) + (1 | task order). This tests the three-way interaction between the categorical fixed effects of ultrasound condition (sham, pulvinar, and central thalamus sonication), the categorical fixed effect of block (pre-sonication and post-sonication), and the continuous fixed effect of minute into the block, which allows us to quantify how each outcome changed after sonication on average as well as over the duration of the task. Descriptive statistics for the cleaned, unaggregated PVT data used in the model are presented in Fig. 2. Estimated marginal means contrasts were used to make more specific comparisons in the models and adjusted for multiple comparisons using the Benjamini-Hochberg method [116]. Key results are presented in the main text. Additional results are described in the Supplemental Material.

### Egly-Driver task (EDT)

Mixed-effects modeling was used to examine whether sonication in any of the ultrasound conditions affected visuospatial attention performance during the EDT, specifically accuracy and response time.

### Data cleaning

Main trials with a recorded response time ≤100 ms were considered false starts and discarded. Participants completed 216.07 main trials (SD = 3.93, min = 205, max = 227) and 23.93 catch trials (SD = 3.93, min = 13, max = 35) per EDT block, on average. The spatial cue appeared ipsilateral to the targeted structure in 13.07 (SD = 2.91, min = 6, max = 21) of the catch trials, and contralateral in 10.86 (SD = 2.52, min = 4, max = 17) of the catch trials, on average. Participants were shown 88.14 validly cued ipsilateral visual targets (SD = 6.44, min = 72, max = 101), 87.37 validly cued contralateral targets (SD = 6.67, min = 64, max = 101), 15.24 invalidly cued ipsilateral targets (SD = 3.06, min = 7, max = 23), and 12.19 invalidly cued contralateral targets (SD = 2.80, min = 6, max = 23) per block, on average. To assess the influence of individual participants on model estimates, we computed the Cook’s distance for each participant in the models. Participants with Cook’s distance values substantially greater than the others (>4 × the mean) were considered influential and their data was examined further. Whole sessions from a participant were left out of the mixed-effects models if they heavily influenced the model estimates and their data was unusual. One session from one participant was left out of the model for catch trials. One session from one participant as left out of the model for main trials. See the Supplemental Material for additional information. We are missing button presses during four EDT blocks due to button box or measurement failure. For one participant, we are missing data for their post-sonication EDT during their sham session. For another participant, we are missing button presses for both the pre- and post-sonication EDT of their pulvinar session as well as the pre-sonication EDT during their sham session. All other data from these participants were used in the mixed-effects models to retain as much data as possible but also to improve estimates of the random effects in the models.

### Analysis

Participants could be either correct (‘hits’ on main trials and ‘correct rejections’ on catch trials) or incorrect (‘misses’ on main trials and ‘false alarms’ on catch trials) on an individual trial, making accuracy a binary variable in the unaggregated EDT data. Two mixed-effects logistic regressions were used to examine the effects of sonication in the different ultrasound conditions on accuracy during the EDT, including one for catch trials and another for main trials. A linear mixed-effects models was used for response time, which was a continuous variable in the unaggregated data. We analyzed response time from main trials only.

### Accuracy

determine whether the effect of sonication in any of the ultrasound conditions on correct responses on catch trials (‘correct rejections’) during the EDT depended on whether the spatial cue appeared ipsilaterally or contralaterally to the ultrasound target, we fit a mixed-effects logistic regression testing the three-way interaction between ultrasound condition (sham, pulvinar, and central thalamus sonication), block (before and after sonication), and spatial cue visual field (ipsilateral or contralateral) accuracy: correct responses ~ condition × block × cue location + (1 + condition | participant) + (1 | task order). Estimated marginal means contrasts were used to make more specific comparisons in the models and adjusted for multiple comparisons using the using Benjamini-Hochberg method.

To examine whether sonication in any of the ultrasound conditions affected correct responses on main trials during the EDT (‘hits’), and if this depended on cue validity and the visual field that the visual target appeared in, we fit a mixed-effect logistic regression testing the three-way interaction between the effects of ultrasound condition (sham, pulvinar, central thalamus sonication), block (before and after sonication), and cue-target location on accuracy: accuracy ~ condition × block × cue-target location + (1 + condition | participant) + (1 | task order). Cue-target location was a categorical variable with four levels: validly cued ipsilateral targets, validly cued contralateral targets, invalidly cued ipsilateral targets, and invalidly cued contralateral targets (see the Supplemental Material Fig. 9). Estimated marginal means contrasts were used to make more specific comparisons in the models and adjusted for multiple comparisons using the using the False Discovery Rate (FDR).

### Response time

To examine whether sonication in any of the ultrasound conditions affected response times on main trials, and if this depended on the spatial cue validity and which visual field the visual target appeared in, we fit a linear mixed-effects model testing three-way interaction between the effects of ultrasound condition (sham, pulvinar, central thalamus sonication), block (before and after sonication), and cue-target location on response time: response time ~ condition × block × cue-target location + (1 + condition | participant) + (1 | task order). Cue-target location was a categorical variable with four levels: validly cued ipsilateral targets, validly cued contralateral targets, invalidly cued ipsilateral targets, and invalidly cued contralateral targets (see the Supplemental Material Fig. 9 for a visualization). Estimated marginal means contrasts were used to make more specific comparisons in the models and adjusted for multiple comparisons using the False Discovery Rate (FDR).

## Acknowledgements

This work was made possible by funding from the Tiny Blue Dot Foundation.

## Author contributions

A.R.H.: Methodology, Software, Validation, Formal Analysis, Investigation, Data Curation, Writing – Original Draft, Writing – Reviewing & Editing, Visualization, Supervision, Project Administration. J.A.C.: Conceptualization, Methodology, Software, Investigation, Data Curation, Supervision, Project Administration. E.A.W.: Investigation, Data Curation, Writing – Original Draft, Writing - Review & Editing. S.M.A.: Investigation, Data Curation, Writing - Review & Editing. M.M.M.: Conceptualization, Methodology, Resources, Writing – Review & Editing, Supervision, Project Administration, Funding Acquisition.

