## Supplemental Material for "Behavioral effects of targeting the central thalamus and pulvinar with transcranial ultrasound stimulation in healthy volunteers"

### Psychomotor Vigilance Task (PVT)

#### *Data cleaning*

Participants completed 210.59 trials (SD = 5.29, min = 197, max = 226) per block, on average. Individual trials were discarded if the response time recorded on that trial was a false start (≤100 ms) or a definite outlier compared to the other trials for that task block and participant based on Tukey's method (less than the first quartile minus 3 × the interquartile range (IQR) or greater than the third quartile plus 3 × the IQR) [1, 2]. There were 6.92 false starts per block per participant, on average (SD = 4.72, min = 1, max = 25). We discarded 6.06 trials for outlier response times per block per participant, on average (SD = 3.55, min = 1, max = 17). We were left with 197.42 trials per block for each participant after trial cleaning, on average (SD = 7.33, min = 176, max = 214).

**Table 1: Trial cleaning results for each participant**

| Participant | Kept (total) | Outlier (total) | Kept (block) | Outlier (block) |
|---|---|---|---|---|
| ***1*** | 1122 | 28 | 187 +/- 7.43 | 4.67 +/- 3.01 |
| ***2*** | 1201 | 30 | 200.16 +/- 4.12 | 5 +/- 2.19 |
| ***3*** | 1187 | 49 | 197.83 +/- 6.49 | 8.17 +/- 2.04 |
| ***4*** | 1141 | 27 | 190.16 +/- 6.59 | 4.50 +/- 3.27 |
| ***5*** | 1243 | 11 | 207.16 +/- 3.19 | 1.83 +/- 0.75 |
| ***6*** | 1206 | 41 | 201 +/- 8.25 | 6.83 +/- 3.11 |
| ***7*** | 1177 | 26 | 167.167 +/- 7.14 | 4.33 +/- 1.97 |
| ***8*** | 1179 | 36 | 196.5 +/- 5.75 | 6 +/- 3.35 |
| ***9*** | 1223 | 12 | 203.83 +/- 5.49 | 2 +/- 1.10 |
| ***10*** | 1181 | 21 | 196.83 +/- 3.92 | 3.5 +/- 1.05 |
| ***11*** | 1161 | 46 | 193.5 +/- 5.96 | 7.66 +/- 2.73 |
| ***12*** | 1194 | 39 | 199 +/- 1.55 | 6.5 +/- 2.17 |
| ***13*** | 1229 | 17 | 204.83 +/- 5.12 | 4.25 +/- 1.71 |
| ***14*** | 1168 | 35 | 194.67 +/- 4.68 | 5.83 +/- 1.94 |
| ***15*** | 1172 | 25 | 195.33 +/- 5.68 | 4.16 +/- 2.40 |
| ***16*** | 1228 | 38 | 204.66 +/- 4.93 | 6.33 +/- 2.25 |
| ***17*** | 1201 | 23 | 200.16 +/- 5.85 | 4.6 +/- 2.19 |
| ***18*** | 1203 | 34 | 200.5 +/-3.45 | 5.67 +/- 2.88 |
| ***19*** | 1209 | 34 | 201.5 +/- 5.47 | 5.67 +/- 2.87 |
| ***20*** | 1220 | 27 | 203.33 +/- 6.47 | 4.5 +/- 2.43 |
| ***21*** | 1130 | 39 | 188.33 +/- 6.38 | 6.5 +/- 2.74 |
| ***22*** | 1140 | 71 | 190 +/- 10.30 | 11.83 +/- 4.18 |
| ***23*** | 1166 | 38 | 194.33 +/- 4.89 | 6.33 +/- 4.23 |
| ***24*** | 1171 | 39 | 195.17 +/- 5 | 6.5 +/- 3.27 |

| | | | | |
|---|---|---|---|---|
| ***25*** | 1195 | 42 | 199.17 +/- 5.78 | 7 +/- 3.85 |
| ***26*** | 1178 | 75 | 196.33 +/- 3.5 | 12.5 +/- 4.18 |
| ***27*** | 1157 | 60 | 192.83 +/- 4.49 | 10 +/- 3.35 |
| ***Avg*** | ***1184.52 +/- 21.22*** | ***35.67 +/- 15.36*** | ***197.42 +/- 7.33*** | ***6.06 +/- 3.55*** |

#### *Additional mixed-effects modeling information and results*

Mixed-effects models were used to test whether sonication in any of the conditions affected response time, response speed, slowest 10% of responses, or lapses during the PVT. Separate models were fit for each outcome but they shared the following specification: outcome ~ condition × block × time into the task block + (1 + condition | participant) + (1 | task order). This tests the three-way interaction between the effects of condition, block, and time into the task on the outcome, as well as all the underlying interactions and main effects. It includes categorical fixed effects for ultrasound condition (sham, pulvinar, and central thalamus sonication) and block (pre- and post-sonication) as well as a continuous fixed effect for time into the task (in minutes). A random intercept for participant with a random slope for condition was added to account for individual differences in the outcome between sessions before sonication, and for having multiple observations per participant since we use unaggregated data from a repeated-measures design. A random intercept for task order was included to account for any differences in the outcomes that can be attributed to the order in which participants completed the tasks. Model summaries are provided for response time (Table 2 and Fig. 2), response speed (Table 4 and Fig. 4), the slowest 10% of response times (Table 6 and Fig. 6), and lapses (Table 8 and Fig. 8). Follow-up estimated marginal means contrasts were used to make more specific comparisons in the models and adjusted for multiple comparisons using the Benjamini-Hochberg method [3].

**Response time.** One participant surpassed the Cook's distance threshold (see Fig. 1A). Upon further examination, this participant showed an erroneous change in response time across blocks during their central thalamus session (see Fig. 1B), which could indicate button box malfunction or another kind of measurement error. We excluded that session for that participant from the mixed-effect model for response time but used all other data in the model. All estimated marginal means contrasts performed are presented in Table 3. Participants showed some variation in their baseline (pre-sonication) response times between the sessions, as shown in the random slopes for ultrasound condition for each participant (see Fig. 2B, right, and Table 2). There was no meaningful variability in response time based on task order (see Fig. 2C and Table 2), indicating that the order in which participants completed the behavioral tasks in did not explain variations in response times. Participants showed the canonical time-on-task effect for response time at baseline (when block is held at pre-sonication and condition is held at sham) whereby response times increased over the duration of the task. Specifically, response times increased by 1.64 ms per minute in the task on average (SE = 6.22, 95% CI = [1.17 – 2.11], $t = 6.81$, $p < 0.001$), as indicated by the fixed effect for minute ('Min') in the model (see Fig. 2A and Table 2). There were no meaningful differences in the average change in response time per minute into the task in any of the conditions or between the conditions, as indicated by the fixed effect terms for the three-way interaction in the model (‘Min:Po-Pr:Pul-Sha’ and ‘Min:Po-Pr:Pul-Sha’ in Table 2) and the estimated marginal means contrasts (see Table 3).

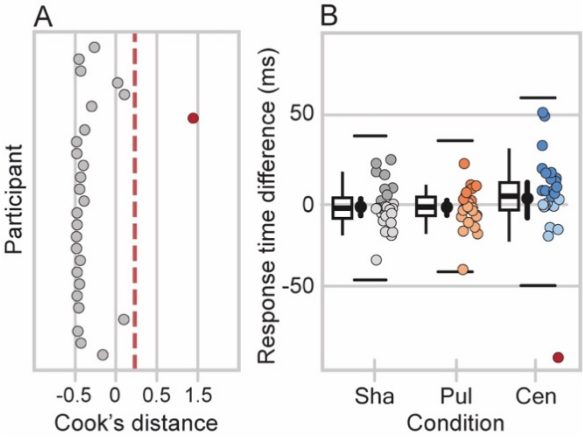


***Supplemental Figure 1:*** *Participant exclusion for the mixed-effects model examining response times during the Psychomotor Vigilance Task (PVT). (A): Cook’s distance for each participant. Red dashed line marks the threshold for exclusion (4 x the mean distance across the participants). One participant (in red) surpassed the Cook’s distance cutoff for exclusion. (B): Distribution of differences in response time between after and before sonication for each condition. Individual points are participants and are lighter or darker if the participant showed greater response times before or after sonication, respectively. One participant (in red) was an outlier in their response time change from pre- to post-sonication during the central thalamus condition. Boxes show first quartile, median, and third quartile and whiskers span the datapoints within 1.5 × the IQR from the lower and upper hinges. Black points and bars show inside each box show mean across participants and 95% confidence intervals. Individual points are participant*

*means. Solid black lines show outer fences based on Tukey's method (less than the first quartile minus 3 × the IQR or greater than the third quartile plus 3 × the IQR.*

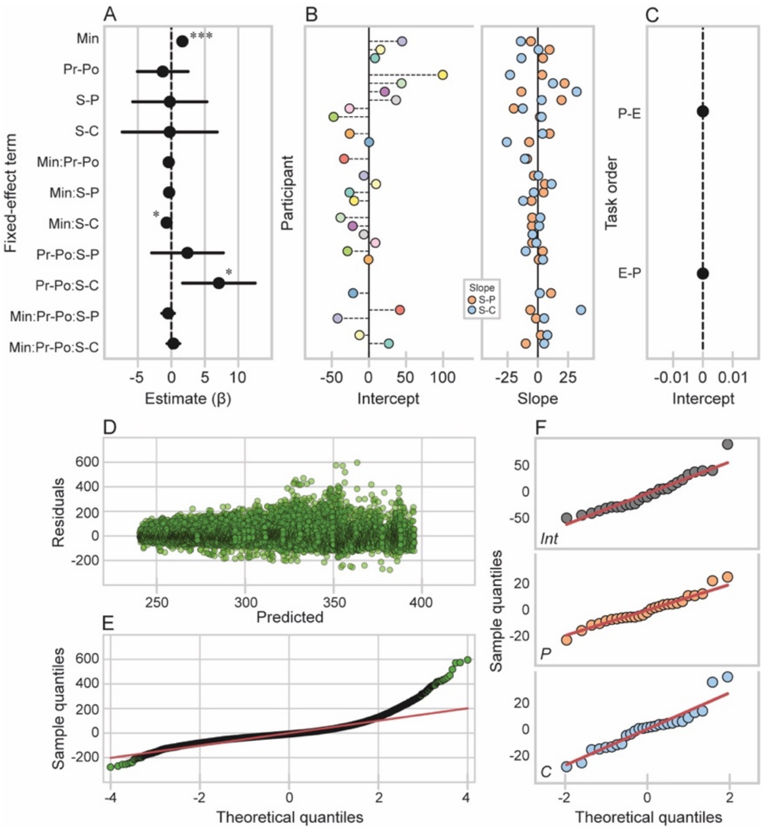


***Supplemental Figure 2:*** *Response time linear mixed-effect model for the Psychomotor Vigilance Task (PVT). (A): Fixed effect estimates. Bars show 95% confidence intervals. (B & C): Random effect estimates. (B): Random intercept for participant (left) with a random slope for ultrasound condition (right). The random intercept for each participant shows their baseline response time (when block is at pre-sonication and ultrasound condition is at sham) relative to the average baseline response time across participants estimated in the fixed effects, which is marked by the black vertical at 0. The random slope for ultrasound condition for each participant shows the difference in their baseline response time (when block is at pre-sonication) between the pulvinar and sham sonication conditions (orange) as well as between the central thalamus and sham sonication conditions (blue) relative to the average across participants estimated in the fixed effect, which is marked by the black vertical line at 0. (C): Random intercept for task order. Points show the average baseline response time (when block is at pre-sonication and ultrasound condition is at sham) for each task order group, specifically PVT then EDT (P-E) and EDT then PVT (E-P). (D): Predicted values versus residuals. (E): QQ plot for theoretical quantiles versus residual (sample) quantiles. (F): QQ plot showing theoretical quantiles versus sample quantiles for the random intercept for participant (top) as well as the random slopes for the steps in condition from sham to pulvinar sonication (middle) and from sham to central thalamus sonication (bottom).*

### Table 2: Linear mixed-effect model: response time ~ minute × block × condition + (1 + condition | participant) + (1 | task order)

**Model summary**

| | *REML* | *Marg $R^2$* | *Cond $R^2$* |
|---|---|---|---|
| | 331620 | 0.004 | 0.32 |

| ***Fit (ANOVA)*** | *AIC* | *BIC* | *Log likelihood* | *Deviance* | $\chi^2$ | *df* | *p* |
|---|---|---|---|---|---|---|---|
| Null/restricted | 338074 | 338150 | -169028 | 338056 | | | |
| Full | 337912 | 338080 | -168936 | 337872 | 184.04 | 11 | **<0.001** |
| ***Fixed effect*** | *Estimate (ms)* | *SE* | *95% CI* | | *t* | *p* | |
| Int | 286.53 | 6.22 | 274.34 – 298.72 | | 46.07 | **<0.001** | |
| Min | 1.64 | 0.24 | 1.17 – 2.11 | | 6.81 | **<0.001** | |
| Pr-Po | -1.29 | 1.93 | -5.07 – 2.50 | | -0.67 | 0.506 | |
| Sha-Pul | -0.23 | 2.83 | -5.78 – 5.32 | | -0.08 | 0.935 | |
| Sha-Cen | -0.24 | 3.62 | -7.34 – 6.85 | | -0.07 | 0.947 | |
| Min:Pr-Po | -0.39 | 0.34 | -1.06 – 0.27 | | -1.16 | 0.246 | |
| Min:Sha-Pul | -0.33 | 0.34 | -1.00 – 0.33 | | -0.98 | 0.327 | |
| Min:Sha-Cen | -0.71 | 0.34 | -1.38 – -0.04 | | -2.07 | **0.038** | |
| Pr-Po:Sha-Pul | 2.40 | 2.74 | -2.96 – 7.76 | | 0.88 | 0.381 | |
| Pr-Po:Cen-Pul | 7.09 | 2.76 | 1.67 – 12.51 | | 2.57 | **0.010** | |
| Min:Po-Pr: Pr-Po:Sha-Pul | -0.48 | 0.48 | -1.43 – 0.46 | | -1.01 | 0.313 | |
| Min:Po-Pr: Pr-Po:Sha-Cen | 0.30 | 0.49 | -0.65 – 1.25 | | 0.62 | 0.537 | |
| ***Random effect*** | *N (obs)* | *Factor* | *Variance* | | *Std* | *Corr* | |
| Participant | 27 (31598) | Int | 993.20 | | 31.52 | | |
| | | Sha-Pul | 114.90 | | 10.72 | 0.17 | |
| | | Sha-Cen | 241.00 | | 15.52 | 0.04 | 0.10 |
| Task order | 2 | Int | 0.00 | | 0.00 | | |
| Residual | | | 2554.20 | | 50.54 | | |

** Model fit with REML. Units in ms. Abbreviations: Marg, marginal; Con, conditional; Min, minute; Pr, pre-sonication; Po, post-sonication; Sha, sham sonication; Pul, pulvinar sonication; Cen, central thalamic sonication. Interaction terms include an ':' and represent the additional change in the included steps across levels of the variable.*

### Table 3: Response time estimated marginal means contrasts

| **Contrast** | **Estimate** | **SE** | **CI (lower)** | **CI (upper)** | **z** | ***p*** | $p_{adj}$ |
|---|---|---|---|---|---|---|---|
| Sha \| Po-Pr | -3.22 | 0.98 | -5.13 | -1.30 | -3.29 | **0.001** | **0.001** |
| Pul \| Po-Pr | -3.20 | 0.98 | -5.12 | -1.28 | -3.27 | **0.001** | **0.001** |
| Cen \| Po-Pr | 5.34 | 1.00 | 3.38 | 7.30 | 5.35 | **<0.001** | **<0.001** |
| Pul-Sha \| Po-Pr | 0.02 | 1.38 | -2.69 | 2.73 | 0.01 | 0.990 | 0.990 |
| Cen-Sha \| Po-Pr | 8.56 | 1.40 | 5.82 | 11.30 | 6.13 | **<0.001** | **<0.001** |
| Cen-Pul \| Po-Pr | 8.55 | 1.40 | 5.80 | 11.29 | 6.11 | **<0.001** | **<0.001** |
| Min (0) \| Sha \| Po-Pr | -1.29 | 1.93 | -5.07 | 2.50 | -0.67 | 0.506 | 0.647 |
| Min (0) \| Pul \| Po-Pr | 1.11 | 1.94 | -2.68 | 4.91 | 0.57 | 0.566 | 0.647 |
| Min (0) \| Cen \| Po-Pr | 5.80 | 1.98 | 1.93 | 9.68 | 2.94 | **0.003** | **0.024** |
| Min (0) \| Cen-Pul \| Po-Pr | 4.69 | 2.77 | -0.73 | 10.11 | 1.70 | 0.090 | 0.212 |
| Min (slope) \| Sha \| Po-Pr | -0.39 | 0.34 | -1.06 | 0.27 | -1.16 | 0.246 | 0.394 |
| Min (slope) \| Pul \| Po-Pr | -0.88 | 0.34 | -1.54 | -0.21 | -2.58 | **0.010** | **0.040** |
| Min (slope) \| Cen \| Po-Pr | -0.09 | 0.35 | -0.77 | 0.59 | -0.27 | 0.787 | 0.787 |
| Min (slope) \| Cen-Pul \| Po-Pr | 0.78 | 0.49 | -0.17 | 1.74 | 1.62 | 0.106 | 0.212 |

**Units in ms. Min (slope) represents the average change per minute into the task. Min (0) represents 0 minutes into the task considered task onset. Abbreviations: Min, minute; Pr, pre-sonication; Po, post-sonication; Sha, sham sonication; Pul, pulvinar sonication; Cen, central thalamic sonication.*

***Response speed.*** The same participant who violated the Cook's criterion for the response time model also surpassed the Cook's distance threshold for response speed (see Fig. 2A and B). We excluded that session for that participant from the mixed-effect model for response speed but used all other data in the model. All estimated marginal means contrasts performed are presented in Table 5. Participants showed some variation in their baseline (pre-sonication) response speeds between the sessions, as shown in the random slopes for ultrasound condition for each participant (see Fig. 4B, right, and Table 4). There was no meaningful variability in response speed based on task order (see Fig. 4C and Table 4). Participants showed the canonical time-on-task effect for response speed at baseline (when block is held at pre-sonication and condition is held at sham) whereby response speeds decreased over the duration of the task. Specifically, response speeds decreased by 0.02 ms per minute in the task on average (SE = 0.00, 95% CI = [-0.02 – -0.01], t = -7.79, p < 0.001, as indicated by the fixed effect for minute ('Min') in the model (see Fig. 4A and Table 4). There were no differences in the average change in response speed per minute after sonication for any condition or between any of the conditions, as indicated by the fixed effect terms for the three-way interaction in the model ('Min:Po-Pr:Pul-Sha' and 'Min:Po-Pr:Pul-Sha' in Table 4) and the estimated marginal means contrasts (see Table 5).

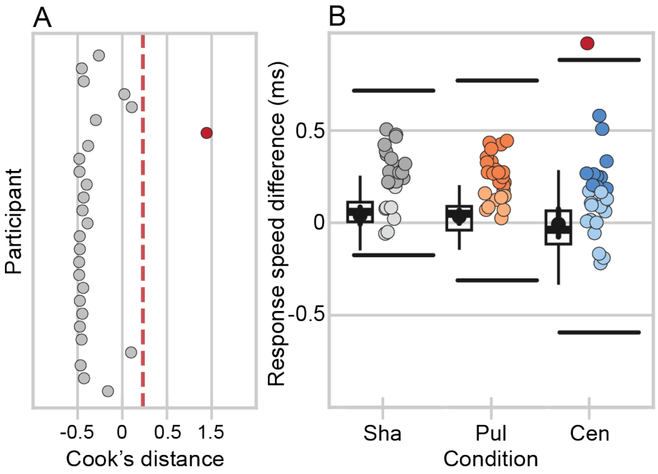


***Supplemental Figure 3:*** *Participant exclusion for the mixed-effects model examining response speeds during the Psychomotor Vigilance Task (PVT). (A): Cook's distance for each participant. Red dashed line marks the threshold for exclusion (4 x the mean distance across the participants). One participant (in red) surpassed the Cook's distance cutoff for exclusion. (B): Distribution of differences in response speed between after and before sonication for each condition. Individual points are participants and are lighter or darker if the participant showed greater response speeds before or after sonication, respectively. One participant (in red) was an outlier in their response speed change from pre- to post-sonication during the central thalamus condition. Boxes show first quartile, median, and third quartile and whiskers span the datapoints within 1.5 × the IQR from the lower and upper hinges. Black points and bars show inside each box show mean across participants and 95% confidence intervals. Individual points are participant*

*means. Solid black lines show outer fences based on Tukey’s method (less than the first quartile minus 3 × the IQR or greater than the third quartile plus 3 × the IQR.*

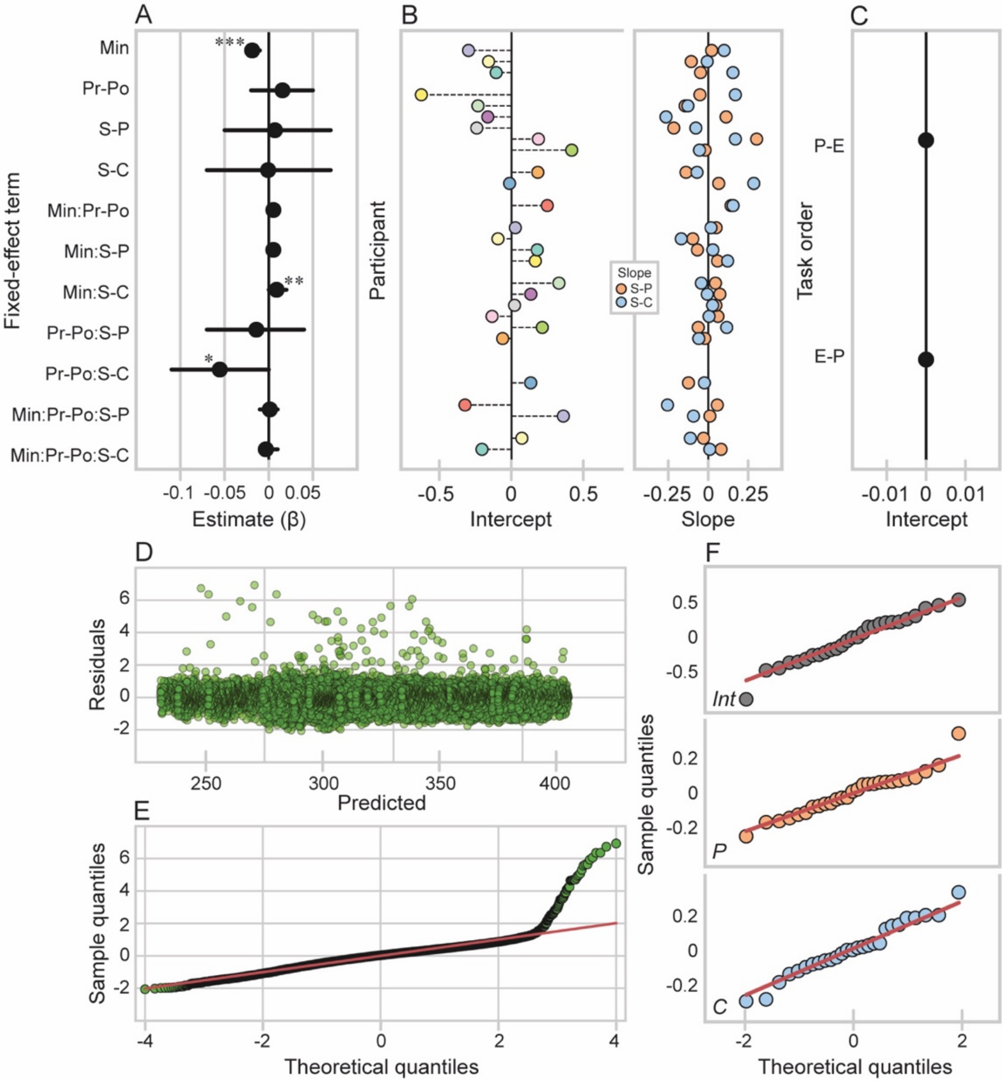


***Supplemental Figure 4:*** *Response speed linear mixed-effect model for the Psychomotor Vigilance Task (PVT). (A): Fixed effect estimates. Bars show 95% confidence intervals. (B & C): Random effect estimates. (B): Random intercept for participant (left) with a random slope for ultrasound condition (right). The random intercept for each participant shows their baseline response speed (when block is at pre-sonication and ultrasound condition is at sham) relative to the average baseline response speed across participants estimated in the fixed effects, which is marked by the black vertical at 0. The random slope for ultrasound condition for each participant shows the difference in their baseline response speed (when block is at pre-sonication) between the pulvinar and sham sonication conditions (orange) as well as between the central thalamus and sham sonication conditions (blue) relative to the average across participants estimated in the fixed effect, which is marked by the black vertical line at 0. (C): Random intercept for task order. Points show the average baseline response speed (when block is at pre-sonication and ultrasound condition is at sham) for each task order group, specifically PVT then EDT (P-E) and EDT then PVT (E-P). (D): Predicted values versus residuals. (E): QQ plot for theoretical quantiles versus residual (sample) quantiles. (F): QQ plot showing theoretical quantiles versus sample quantiles for the random intercept for participant (top) as well as the random slopes for the steps in condition from sham to pulvinar sonication (middle) and from sham to central thalamus sonication (bottom).*

**Table 4: Linear mixed-effect model: response speed ~ minute × block × condition + (1 + condition | participant) + (1 | task order)**

**Model summary**

| | | | | | | | |
|---|---|---|---|---|---|---|---|
| | *REML* | *Marg $R^2$* | *Cond $R^2$* | | | | |
| | 46857 | 0.005 | 0.32 | | | | |
| ***Fit (ANOVA)*** | *AIC* | *BIC* | *Log likelihood* | *deviance* | $\chi^2$ | *df* | *p* |
| Null/restricted | 46979 | 47054 | -23480 | 46961 | | | |
| Full | 46799 | 46966 | -23380 | 46759 | 201.48 | 11 | **<0.001** |
| ***Fixed effect*** | *Estimate (ms)* | *SE* | *95% CI* | *t* | *p* | | |
| Int | 3.60 | 0.06 | 3.48 – 3.72 | 57.62 | **<0.001** | | |
| Min | -0.02 | 0.00 | -0.02 – -0.01 | -7.79 | **<0.001** | | |
| Pr-Po | 0.02 | 0.02 | -0.02 – 0.05 | 0.80 | 0.422 | | |
| Sha-Pul | 0.01 | 0.03 | -0.05 – 0.07 | 0.23 | 0.819 | | |
| Sha-Cen | -0.00 | 0.04 | -0.07 – 0.07 | -0.02 | 0.982 | | |
| Min:Pr-Po | 0.01 | 0.00 | -0.00 – 0.01 | 1.54 | 0.124 | | |
| Min:Sha-Pul | 0.01 | 0.00 | -0.00 – 0.01 | 1.52 | 0.128 | | |
| Min:Sha-Cen | 0.01 | 0.00 | 0.00 – 0.02 | 2.57 | **0.010** | | |
| Pr-Po:Sha-Pul | -0.01 | 0.03 | -0.07 – 0.04 | -0.52 | 0.606 | | |
| Pr-Po:Cen-Pul | -0.06 | 0.03 | -0.11 – -0.00 | -2.01 | **0.045** | | |
| Min:Po-Pr: Pr-Po:Sha-Pul | 0.00 | 0.00 | -0.01 – 0.01 | 0.24 | 0.809 | | |
| Min:Po-Pr: Pr-Po:Sha-Cen | -0.00 | 0.00 | -0.01 – 0.01 | -0.67 | 0.501 | | |
| ***Random effect*** | *N (obs)* | *Factor* | *Variance* | *Std* | *Corr* | | |
| Participant | 27 (31598) | Int | 0.100 | 0.317 | | | |
| | | Sha-Pul | 0.015 | 0.121 | 0.18 | | |
| | | Sha-Cen | 0.022 | 0.149 | 0.06 | 0.29 | |
| Task order | 2 | Int | 0.000 | 0.000 | | | |
| Residual | | | 0.255 | 0.505 | | | |

** Model fit with REML. Units in ms. Abbreviations: Marg, marginal; Con, conditional; Min, minute; Pr, pre-sonication; Po, post-sonication; Sha, sham sonication; Pul, pulvinar sonication; Cen, central thalamic sonication. Interaction terms include an ':' and represent the additional change in the included steps across levels of the variable.*

**Table 5: Response speed estimated marginal means contrasts**

| Contrast | Estimate | SE | CI (lower) | CI (upper) | z | *p* | $p_{adj}$ |
|---|---|---|---|---|---|---|---|
| Sha \| Po-Pr | 0.04 | 0.01 | 0.02 | 0.06 | 4.21 | **<0.001** | **<0.001** |
| Pul \| Po-Pr | 0.03 | 0.01 | 0.01 | 0.05 | 3.35 | **0.001** | **0.002** |
| Cen \| Po-Pr | -0.03 | 0.01 | -0.05 | -0.01 | -3.04 | **0.002** | **0.002** |
| Pul-Sha \| Po-Pr | -0.01 | 0.01 | -0.04 | 0.02 | -0.61 | 0.545 | 0.545 |
| Cen-Sha \| Po-Pr | -0.07 | 0.01 | -0.10 | -0.04 | -5.12 | **0.000** | **0.000** |
| Cen-Pul \| Po-Pr | -0.06 | 0.01 | -0.09 | -0.04 | -4.51 | **0.000** | **0.000** |
| Min (0) \| Sha \| Po-Pr | 0.02 | 0.02 | -0.02 | 0.05 | 0.80 | 0.422 | 0.563 |
| Min (0) \| Pul \| Po-Pr | 0.00 | 0.02 | -0.04 | 0.04 | 0.07 | 0.941 | 0.941 |
| Min (0) \| Cen \| Po-Pr | -0.04 | 0.02 | -0.08 | 0.00 | -2.02 | **0.043** | 0.240 |
| Min (0) \| Cen-Pul \| Po-Pr | -0.04 | 0.03 | -0.10 | 0.01 | -1.50 | 0.135 | 0.270 |
| Min (slope) \| Sha \| Po-Pr | 0.01 | 0.00 | 0.00 | 0.01 | 1.54 | 0.124 | 0.270 |
| Min (slope) \| Pul \| Po-Pr | 0.01 | 0.00 | 0.00 | 0.01 | 1.88 | 0.060 | 0.240 |

| | | | | | | | |
|---|---|---|---|---|---|---|---|
| Min (slope) \| Cen \| Po-Pr | 0.00 | 0.00 | 0.00 | 0.01 | 0.56 | 0.573 | 0.655 |
| Min (slope) \| Cen-Pul \| Po-Pr | 0.00 | 0.00 | -0.01 | 0.01 | -0.91 | 0.362 | 0.563 |

**Units in ms. Min (slope) represents the average change per minute into the task. Min (0) represents 0 minutes into the task considered task onset. Abbreviations: Min, minute; Pr, pre-sonication; Po, post-sonication; Sha, sham sonication; Pul, pulvinar sonication; Cen, central thalamic sonication.*

**Slowest 10% of responses*.*** Three participants surpassed the Cook's distance threshold (see Fig. 5A). This included the same participant as the one listed for response time and response speed. All these participants showed erroneous changes in the slowest 10% of their responses across blocks during at least one of the sonication sessions (see Fig. 5B), which could indicate button box malfunction or greater variability in the slowest responses. We excluded these sessions from the mixed-effects model for the slowest 10% of response time but kept all other sessions from these participants in the model. All estimated marginal means contrasts performed are presented in Table 7. Participants showed some variation in their baseline (pre-sonication) response speeds between the sessions, as shown in the random slopes for ultrasound condition for each participant (see Fig. 6B, right, and Table 6). There was some variability in response speed based on task order (see Fig. 6C and Table 6), indicating that the random effect accounted for some variations in the slowest responses attributable to the order in which participants completed the behavioral tasks.

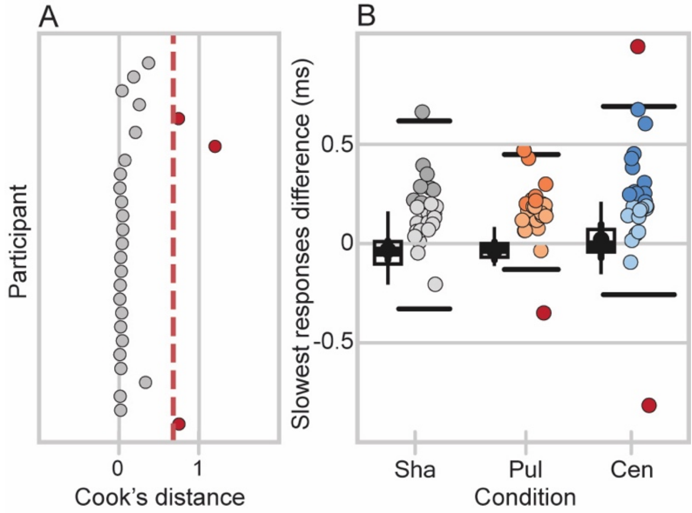


***Supplemental Figure 5:*** *Participant exclusion for the mixed-effects model examining the slowest 10% of responses during the Psychomotor Vigilance Task (PVT). (A): Cook's distance for each participant. Red dashed line marks the threshold for exclusion (4 x the mean distance across the participants). One participant (in red) surpassed the Cook's distance cutoff for exclusion. (B): Distribution of differences in response time between after and before sonication for each condition. Individual points are participants and are lighter or darker if the participant showed greater response times before or after sonication, respectively. One participant (in red) was an outlier in their response time change from pre- to post-sonication during the central thalamus condition. Boxes show first quartile, median, and third quartile and whiskers span the datapoints within 1.5 × the IQR from the lower and upper hinges. Black points and bars show inside each box show mean across participants and 95% confidence intervals. Individual points are*

*participant means. Solid black lines show outer fences based on Tukey's method (less than the first quartile minus 3 × the IQR or greater than the third quartile plus 3 × the IQR.*

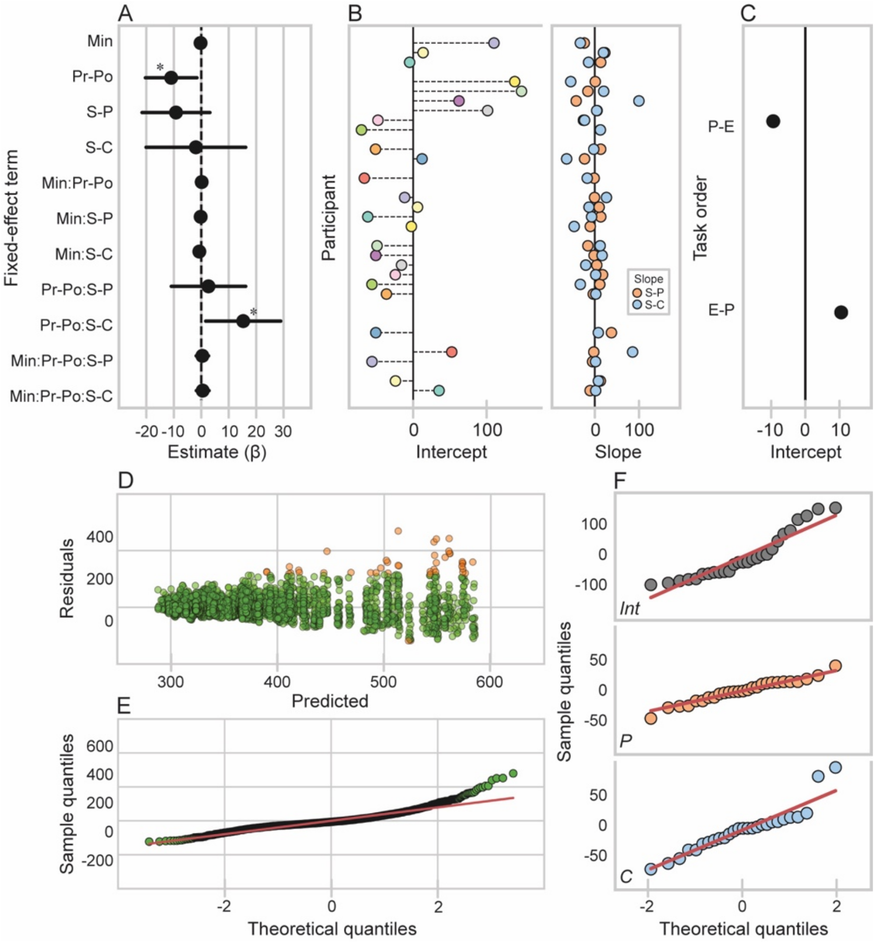


***Supplemental Figure 6:*** *Slowest 10% of responses linear mixed-effect model for the Psychomotor Vigilance Task (PVT). (A): Fixed effect estimates. Bars show 95% confidence intervals. (B & C): Random effect estimates. (B): Random intercept for participant (left) with a random slope for ultrasound condition (right). The random intercept for each participant shows their baseline response time (when block is at pre-sonication and ultrasound condition is at sham) relative to the average baseline response time across participants estimated in the fixed effects, which is marked by the black vertical at 0. The random slope for ultrasound condition for each participant shows the difference in their baseline response time (when block is at pre-sonication) between the pulvinar and sham sonication conditions (orange) as well as between the central thalamus and sham sonication conditions (blue) relative to the average across participants estimated in the fixed effect, which is marked by the black vertical line at 0. (C): Random intercept for task order. Points show the average baseline response time (when block is at pre-sonication and ultrasound condition is at sham) for each task order group, specifically PVT then EDT (P-E) and EDT then PVT (E-P). (D): Predicted values versus residuals. (E): QQ plot for theoretical quantiles versus residual (sample) quantiles. (F): QQ plot showing theoretical quantiles versus sample quantiles for the random intercept for participant (top) as well as the random slopes for the steps in condition from sham to pulvinar sonication (middle) and from sham to central thalamus sonication (bottom).*

### Table 6: Linear mixed-effect model: slowest 10% of responses ~ minute × block × condition + (1 + condition | participant) + (1 | task order)

**Model summary**

| | | | | | | | |
|---|---|---|---|---|---|---|---|
| | *REML* | *Marg $R^2$* | *Cond $R^2$* | | | | |
| | 32221 | 0.005 | 0.81 | | | | |
| ***Fit (ANOVA)*** | *AIC* | *BIC* | *Log likelihood* | *Deviance* | $\chi^2$ | *df* | *p* |
| Null/restricted | 32313 | 32367 | -16148 | 32295 | | | |
| Full | 32294 | 32415 | -16127 | 32254 | 41.11 | 11 | **<0.001** |
| ***Fixed effect*** | *Estimate* | *SE* | *95% CI* | *t* | *p* | | |
| Int | 398.24 | 20.54 | 357.98 – 438.51 | 19.39 | **<0.001** | | |
| Min | -0.25 | 0.57 | -1.37 – 0.87 | -0.43 | 0.664 | | |
| Pr-Po | -10.95 | 4.79 | -20.35 – -1.56 | -2.29 | **0.022** | | |
| Sha-Pul | -9.21 | 6.30 | -21.57 – 3.15 | -1.46 | 0.144 | | |
| Sha-Cen | -1.96 | 9.26 | -20.12 – 16.20 | -0.21 | 0.832 | | |
| Min:Pr-Po | 0.16 | 0.80 | -1.40 – 1.73 | 0.20 | 0.840 | | |
| Min:Sha-Pul | -0.21 | 0.82 | -1.82 – 1.40 | -0.25 | 0.802 | | |
| Min:Sha-Cen | -0.70 | 0.84 | -2.35 – 0.95 | -0.83 | 0.404 | | |
| Pr-Po:Sha-Pul | 2.64 | 6.91 | -10.92 – 16.20 | 0.38 | 0.703 | | |
| Pr-Po:Cen-Pul | 15.30 | 6.97 | 1.62 – 28.97 | 2.19 | **0.028** | | |
| Min:Po-Pr:Pr-Po:Sha-Pul | 0.41 | 1.15 | -1.84 – 2.67 | 0.36 | 0.720 | | |
| Min:Po-Pr:Pr-Po:Sha-Cen | 0.53 | 1.18 | -1.78 – 2.84 | 0.45 | 0.652 | | |
| ***Random effect*** | *N (obs)* | *Factor* | *Variance* | *Std* | *Corr* | | |
| Participant | 27 (3153) | Int | 5418.9 | 73.61 | | | |
| | | Sha-Pul | 386.8 | 19.67 | -0.39 | | |
| | | Sha-Cen | 1519.9 | 38.99 | 0.08 | -0.05 | |
| Task order | 2 | Int | 416.0 | 20.40 | | | |
| Residual | | | 1484.6 | 38.53 | | | |

** Model fit with REML. Units in ms. Abbreviations: Marg, marginal; Con, conditional; Min, minute; Pr, pre-sonication; Po, post-sonication; Sha, sham sonication; Pul, pulvinar sonication; Cen, central thalamic sonication. Interaction terms include an ':' and represent the additional change in the included steps across levels of the variable.*

### Table 7: Slowest 10% estimated marginal means contrasts

| Contrast | Estimate | SE | CI (lower) | CI (upper) | z | *p* | $p_{adj}$ |
|---|---|---|---|---|---|---|---|
| Sha \| Po-Pr | -10.11 | 2.33 | -14.67 | -5.55 | -4.35 | **<0.001** | **<0.001** |
| Pul \| Po-Pr | -5.32 | 2.38 | -9.98 | -0.65 | -2.23 | **0.025** | **0.030** |
| Cen \| Po-Pr | 7.96 | 2.44 | 3.19 | 12.73 | 3.27 | **0.001** | **0.002** |
| Pul-Sha \| Po-Pr | 4.79 | 3.33 | -1.73 | 11.32 | 1.44 | 0.150 | 0.150 |
| Cen-Sha \| Po-Pr | 18.07 | 3.37 | 11.47 | 24.67 | 5.37 | **<0.001** | **<0.001** |
| Cen-Pul \| Po-Pr | 13.28 | 3.40 | 6.60 | 19.95 | 3.90 | **<0.001** | **<0.001** |
| Min (0) \| Sha \| Po-Pr | -10.95 | 4.79 | -20.34 | -1.57 | -2.29 | **0.022** | 0.176 |
| Min (0) \| Pul \| Po-Pr | -8.31 | 4.99 | -18.09 | 1.46 | -1.67 | 0.095 | 0.253 |
| Min (0) \| Cen \| Po-Pr | 4.34 | 5.07 | -5.59 | 14.28 | 0.86 | 0.392 | 0.652 |
| Min (0) \| Cen-Pul \| Po-Pr | 12.66 | 7.11 | -1.28 | 26.59 | 1.78 | 0.075 | 0.253 |
| Min (slope) \| Sha \| Po-Pr | 0.16 | 0.80 | -1.40 | 1.73 | 0.20 | 0.840 | 0.921 |
| Min (slope) \| Pul \| Po-Pr | 0.57 | 0.83 | -1.05 | 2.20 | 0.69 | 0.489 | 0.652 |
| Min (slope) \| Cen \| Po-Pr | 0.69 | 0.86 | -1.00 | 2.39 | 0.80 | 0.424 | 0.652 |
| Min (slope) \| Cen-Pul \| Po-Pr | 0.12 | 1.20 | -2.23 | 2.46 | 0.10 | 0.921 | 0.921 |

**Units in ms. Min (slope) represents the average change per minute into the task. Min (0) represents 0 minutes into the task considered task onset. Abbreviations: Min, minute; Pr, pre-sonication; Po, post-sonication; Sha, sham sonication; Pul, pulvinar sonication; Cen, central thalamic sonication.*

**Lapses.** Lapses refer to lapses in attention and are unusually delayed responses. Response times greater than 500 ms or twice the mean response time for an individual participant are traditionally considered lapses [4]. However, our participants never or rarely showed response times greater than 500 ms or twice their mean response time, even before trial cleaning. Instead, we combined the PVT data from all blocks for each participant and considered lapses to be response times greater than the mean response time plus two standard deviations for that participant (see Supplemental Fig. 7). On average, out of the 1129.37 trials total that each participant completed across all blocks (SD = 32.96, min = 1075, max = 1188), there were 55.15 lapses total (SD = 7.98, min = 40, max = 69).

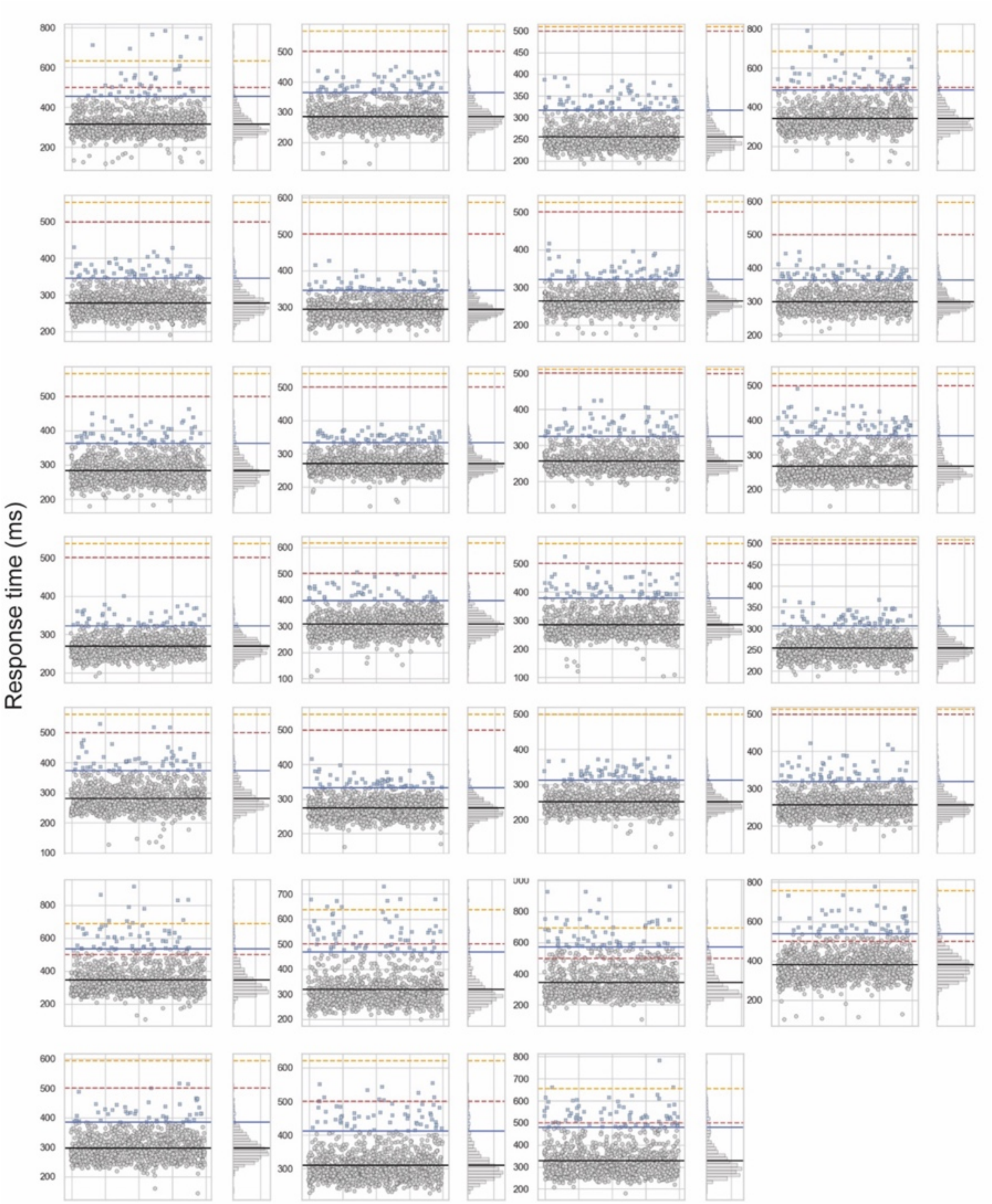


***Supplemental Figure 7:*** *Defining lapses in attention during the Psychomotor Vigilance Task (PVT) for each participant. Each subplot shows the distribution of individual response times (grey points) and lapses in attention (blue plus symbols) for each participant for all blocks. The solid black line shows the grand mean response time for that participant. The solid blue line shows the lapse cutoff for the participant (*their mean response time plus two standard deviations*). The red dashed line shows the 500 ms mark. The yellow dashed line shows twice the mean response time.*

No participants surpassed the Cook’s distance threshold for the lapses model. However, since there was an error in the response time data from one participant’s session (see Fig. 1A) and lapses are derived from response time, we left that session for that participant out of the mixed-effects model for lapses, as well. All other data were used in the model. Participants showed some variation in their baseline (pre-sonication) lapse probability between the ultrasound sessions, as indicated by the random slopes for condition per participant (see Table 8 and Fig. 8B, right). There was no meaningful variability in lapse probability based on task order (see Table 8 and Fig. 8C), indicating that the order in which participants completed the behavioral tasks did not contribute to the variability in lapses. Participants showed the canonical time-on-task effect at baseline (pre-sonication in the sham condition) whereby the frequency of lapses increased over the duration of the task. Specifically, participants were 1.05 times more likely to lapse per minute in the task (95% CI = [1.01, 1.09], $z = 2.34$, $p = 0.002$), as indicated by the fixed effect term for minute in the model (see Fig. 8A and Table 8). All estimated marginal mean contrasts performed for lapses are presented in Table 9.

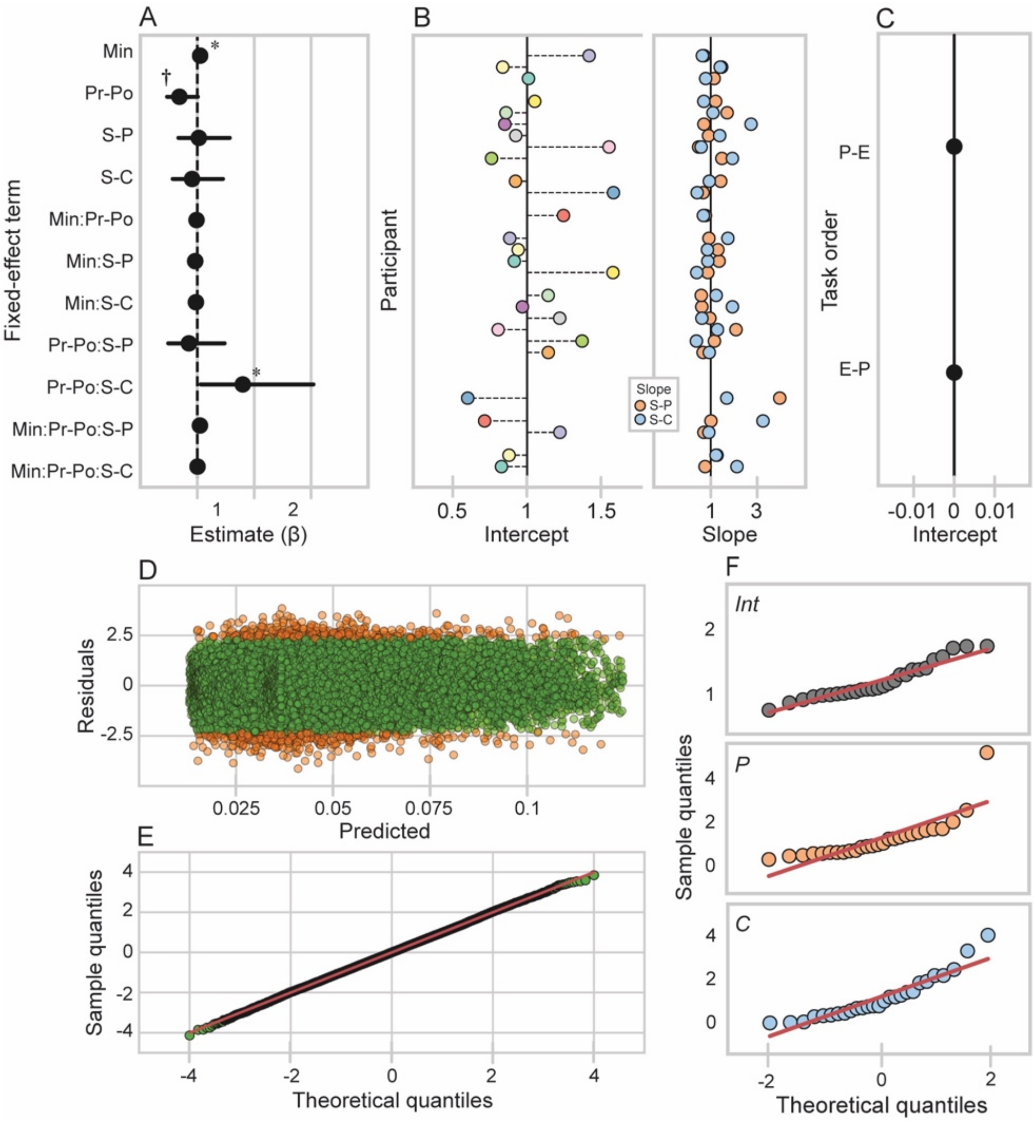

***Supplemental Figure 8:*** *Mixed-effects logistic regression for lapses in attention during the Psychomotor Vigilance Task (PVT). (A): Fixed effect estimates. Bars show 95% confidence intervals. (B & C): Random effect estimates. (B): Random intercept for participant (left) with a random slope for ultrasound condition (right). The random intercept for each participant shows their baseline odds of lapsing (when block is at pre-sonication and ultrasound condition is at sham) relative to the average baseline odds of lapsing across participants estimated in the fixed effects, which is marked by the black vertical at 0. The random slope for ultrasound condition for each participant shows the difference in their baseline odds of lapsing (when block is at pre-sonication) between the pulvinar and sham sonication conditions (orange) as well as between the central thalamus and sham sonication conditions (blue) relative to the average across participants estimated in the fixed effect, which is marked by the black vertical line at 0. (C): Random intercept for task order. Points show the average baseline odds of lapsing (when block is at pre-sonication and ultrasound condition is at sham) for each task order group, specifically PVT then EDT (P-E) and EDT then PVT (E-P). (D): Predicted values (odds ratios) versus simulated (and normalized) residuals. (E): QQ plot for theoretical quantiles versus simulated (and normalized) sample residual quantiles. (F): QQ plot showing theoretical quantiles versus sample quantiles for the random intercept for participant (top) as well as the random slopes for the steps in condition from sham to pulvinar sonication (middle) and from sham to central thalamus sonication (bottom).*

### Table 8: Mixed-effects logistic regression: lapses in attention ~ minute × block × condition + (1 + condition | participant) + (1 | task order)

**Model summary**

| | *Marg $R^2$* | *Cond $R^2$* |
|---|---|---|
| | 0.014 | 0.07 |

| ***Fit (ANOVA)*** | *AIC* | *BIC* | *Log likelihood* | *Deviance* | $\chi^2$ | *df* | *p* |
|---|---|---|---|---|---|---|---|
| Null/restricted | 11657 | 11724 | -5820.6 | 11641 | | | |
| Full | 11625 | 11784 | -5793.5 | 11587 | 54.11 | 11 | **<0.001** |

| ***Fixed effect*** | *Estimate* | *95% CI* | *z* | *p* |
|---|---|---|---|---|
| Int | 0.04 | 0.03 – 0.06 | -21.50 | **<0.001** |
| Min | 1.05 | 1.01 – 1.09 | 2.34 | **0.019** |
| Pr-Po | 0.69 | 0.47 – 1.01 | -1.93 | 0.054 |
| Sha-Pul | 1.03 | 0.67 – 1.57 | 0.12 | 0.902 |
| Sha-Cen | 0.91 | 0.57 – 1.45 | -0.40 | 0.688 |
| Min:Pr-Po | 0.99 | 0.93 – 1.05 | -0.41 | 0.683 |
| Min:Sha-Pul | 0.96 | 0.91 – 1.02 | -1.23 | 0.218 |
| Min:Sha-Cen | 0.98 | 0.92 – 1.04 | -0.78 | 0.433 |
| Pr-Po:Sha-Pul | 0.85 | 0.49 – 1.48 | -0.57 | 0.570 |
| Pr-Po:Cen-Pul | 1.80 | 1.06 – 3.04 | 2.19 | **0.028** |
| Min:Po-Pr:Pr-Po:Sha-Pul | 1.05 | 0.95 – 1.15 | 0.97 | 0.333 |
| Min:Po-Pr:Pr-Po:Sha-Cen | 1.00 | 0.92 – 1.10 | 0.10 | 0.922 |

| ***Random effect*** | *N (obs)* | *Factor* | *Variance* | *Std* | *Corr* | |
|---|---|---|---|---|---|---|
| Participant | 27 (31598) | Int | 1.51 | 1.46 | | |
| | | Sha-Pul | 1.47 | 1.86 | -0.70 | |
| | | Sha-Cen | 1.83 | 2.16 | -0.82 | 0.23 |
| Task order | 2 | Int | 1.00 | 1.00 | | |

** Units in odds ratios. An odds ratio of 1 reflects equal likelihood of lapsing. Abbreviations: Marg, marginal; Con, conditional; Min, minute; Pr, pre-sonication; Po, post-sonication; Sha, sham sonication; Pul, pulvinar sonication; Cen, central thalamic sonication. Interaction terms include an ':' and represent the additional change in the included steps across levels of the variable.*

### Table 9: Lapses estimated marginal means contrasts

| Contrast | Odds ratio | CI (lower) | CI (upper) | z | *p* | $p_{adj}$ |
|---|---|---|---|---|---|---|
| Sha \| Po-Pr | 0.64 | 0.53 | 0.77 | -4.66 | **<0.001** | **<0.001** |
| Pul \| Po-Pr | 0.69 | 0.57 | 0.83 | -3.80 | **<0.001** | **<0.001** |
| Cen \| Po-Pr | 1.18 | 0.99 | 1.41 | 1.84 | 0.066 | 0.079 |
| Pul-Sha \| Po-Pr | 1.07 | 0.82 | 1.40 | 0.48 | 0.630 | 0.630 |
| Cen-Sha \| Po-Pr | 1.84 | 1.42 | 2.38 | 4.64 | **<0.001** | **<0.001** |
| Cen-Pul \| Po-Pr | 1.72 | 1.32 | 2.24 | 4.04 | **<0.001** | **<0.001** |
| Min (0) \| Sha \| Po-Pr | 0.69 | 0.47 | 1.01 | -1.93 | 0.054 | 0.144 |
| Min (0) \| Pul \| Po-Pr | 0.59 | 0.39 | 0.87 | -2.65 | **0.008** | **0.032** |
| Min (0) \| Cen \| Po-Pr | 1.24 | 0.86 | 1.77 | 1.15 | 0.250 | 0.497 |
| Min (0) \| Cen-Pul \| Po-Pr | 2.11 | 1.24 | 3.60 | 2.74 | **0.006** | **0.032** |
| Min (slope) \| Sha \| Po-Pr | 0.99 | 0.93 | 1.05 | -0.41 | 0.683 | 0.777 |
| Min (slope) \| Pul \| Po-Pr | 1.03 | 0.97 | 1.10 | 0.95 | 0.343 | 0.497 |
| Min (slope) \| Cen \| Po-Pr | 0.99 | 0.93 | 1.05 | -0.28 | 0.777 | 0.777 |
| Min (slope) \| Cen-Pul \| Po-Pr | 0.96 | 0.88 | 1.05 | -0.89 | 0.373 | 0.497 |

** Units in odds ratios. An odds ratio of 1 reflects equal likelihood of lapsing. Min (slope) represents the average change per minute into the task. Min (0) represents 0 minutes into the task considered task onset. Abbreviations: Min, minute; Pr, pre-sonication; Po, post-sonication; Sha, sham sonication; Pul, pulvinar sonication; Cen, central thalamic sonication.*

### Edgly-Driver Task (EDT)

#### *Data cleaning*

There were 1.83 false starts (SD = 3.35, min = 0, max = 20) per block, on average. Participants completed 216.07 main trials (SD = 3.93, min = 205, max = 227) and 23.93 catch trials (SD = 3.93, min = 13, max = 35) per EDT block, on average. The spatial cue appeared ipsilateral to the targeted structure in 13.07 (SD = 2.91, min = 6, max = 21) of the catch trials, and contralateral in 10.86 (SD = 2.52, min = 4, max = 17) of the catch trials, on average. We considered all validly cued main trials but limited invalidly cued trials to those where the spatial cue and visual target appeared in different sides of the visual field. This separated the main trials into four groups based on spatial cue validily and visual target location, including trials with vaidly cued ipsilateral visual targets, invalidly cued ipsilateral targets, validlliy cued contralateral targets, and invalidly cued contralateral targets (see Supplemental Fig 9 for a visualization). Participants were shown 88.14 validly cued ipsilateral targets (SD = 6.44, min = 72, max = 101), 87.37 validly cued contralateral targets (SD = 6.67, min = 64, max = 101), 15.24 invalidly cued ipsilateral targets (SD = 3.06, min = 7, max = 23), and 12.19 invalidly cued contralateral targets(SD = 2.80, min = 6, max = 23) per block, on average.

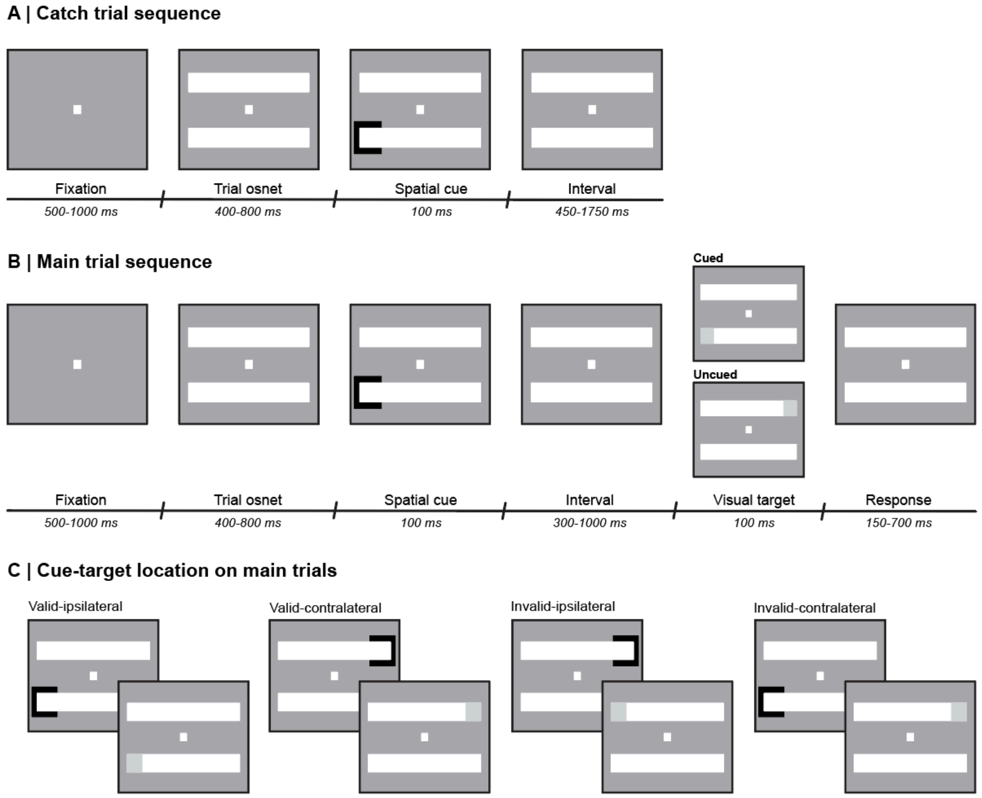


***Supplemental Fig. 9:*** *Schematic of the Edgly-Driver Task (EDT). (A): Catch trial. Spatial cue appears to direct the participant's attention and the trial ends. (B): Main trial. Spatial cue appears to direct the participant's attention to a location and then a visual target appears in either the cued location or another location. (C): Grouping of main trials according to the relationship between spatial cue and visual target location. Stimuli could appear in the ipsilateral or contralateral visual field relative to the sonicated region (left pulvinar and left central thalamus). There were four kinds of main trials based on cue validity and*

*visual target location:* validly cued ipsilateral targets, validly cued contralateral targets, invalidly cued ipsilateral targets, and invalidly cued contralateral targets.

#### ***Additional mixed-effects modeling information and results***

Mixed-effects modeling was used to examine whether sonication in any of the ultrasound conditions affected visuospatial attention during the EDT, specifically correct responses and response time. All models included a random intercept for participant with a random slope of condition (1 + condition | participant) as well as a random intercept for task order (1 | task order). The random intercept for participant with the random slope for condition accounts for individual differences in the outcome between sessions before sonication, and for having multiple observations per participant since we use unaggregated data from a repeated-measures design. The random intercept for task order captures any differences in the outcomes that can be attributed to the order in which participants completed the behavioral tasks.

**Accuracy.** Participants could be either correct (‘hits’ on main trials and ‘correct rejections’ on catch trials) or incorrect (‘misses’ on main trials and ‘false alarms’ on catch trials) on a trial, making accuracy a binary variable in the unaggregated EDT data. Two mixed-effects logistic regressions were thus used to examine the effects of sonication on correct responses during the EDT, including one for catch trials and another for main trials.

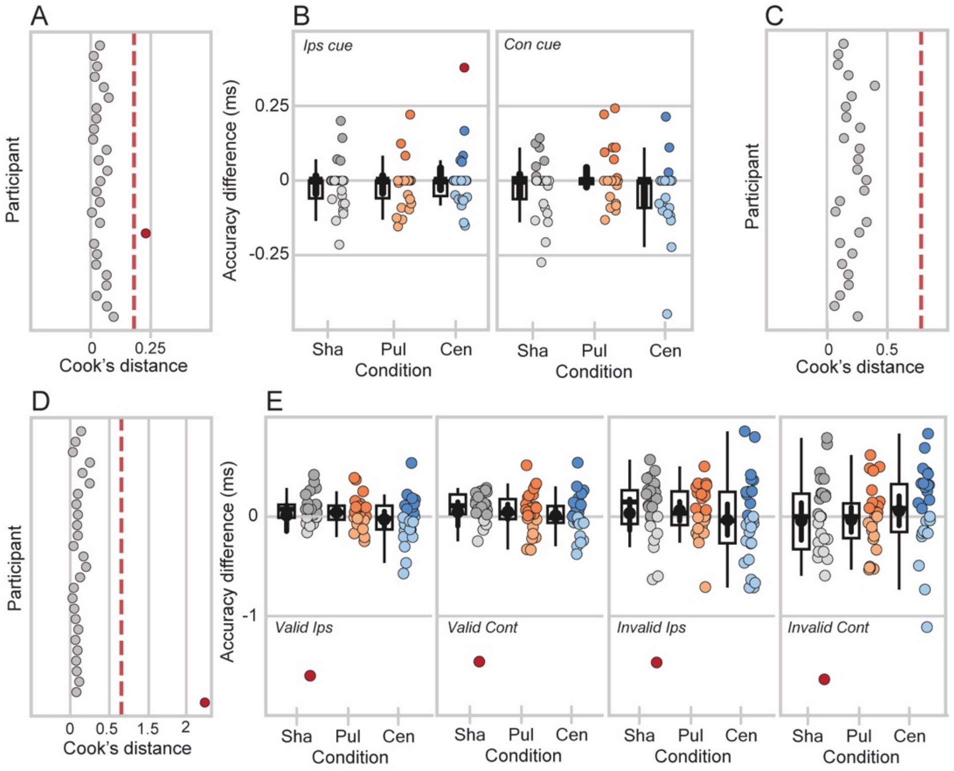


***Supplemental Figure 10:*** *Participant exclusion for the mixed-effects models for the Edgly-Driver Task (EDT). (A & B): Mixed-effects model for catch trial accuracy. (D & E): Mixed-effects model for main trial accuracy. (C): Mixed-effects model for response time. (A, C, & D): Cook’s distance for each participant. Red dashed line marks the threshold for exclusion (4 x the mean distance across the participants).*

*Participants flagged are shown in red. (B & E): Distribution of differences in accuracy between after and before sonication for each condition across the different types of catch trials (B) and main trials (E). Individual points are participants and are lighter or darker if the participant showed greater response times before or after sonication, respectively. Participants flagged based on their Cook's distanced appear red. Boxes show first quartile, median, and third quartile and whiskers span the datapoints within 1.5 × the IQR from the lower and upper hinges. Black points and bars show inside each box show mean across participants and 95% confidence intervals. Individual points are participant means.*

***Catch trials.*** To assess the effects of sonication on correct responses during catch trials ('correct rejections'), we fit a mixed-effects logistic regression testing the three-way interaction between ultrasound condition (sham, pulvinar, and central thalamus sonication), block (before and after sonication), and spatial cue location (ipsilateral or contralateral to the ultrasound target) on accuracy: accuracy ~ condition × block × cue location + (1 + condition | participant) + (1 | task order). One participant surpassed the Cook's distance threshold (see Fig. 10A). Under further evaluation, this participant showed an erroneous change in accuracy across blocks during their central thalamus session (see Fig. 10B). So, we excluded that session for that participant from the mixed-effect model. All other data were used in the model. Participants showed some variation in their baseline (pre-sonication) probability of responding correctly on catch trials ('correct rejections') between the sessions, as indicated by the random slopes for condition for each participant (see Table 10 and Fig. 11B, right). There was no meaningful variability in the probability of responding correctly between the task order groups (see Table 10 and Fig. 11C) captured by the model. All estimated marginal means contrasts performed are presented in Table 11. Follow-up estimated marginal means contrasts were used to make more specific comparisons in the models and adjusted for multiple comparisons using the Benjamini-Hochberg method.

**Table 10: Mixed-effect logistic regression: catch trial accuracy ~ spatial cue location × block × condition + (1 + condition | participant) + (1 | task order)**

**Model summary**

| | | | | | | | |
|---|---|---|---|---|---|---|---|
| | *Marg $R^2$* | *Cond $R^2$* | | | | | |
| | 0.02 | 0.37 | | | | | |
| ***Fit (ANOVA)*** | *AIC* | *BIC* | *Log likelihood* | *Deviance* | $\chi^2$ | *df* | *p* |
| Null/restricted | 1351.3 | 1401.1 | -667.65 | 1335.3 | | | |
| Full | 1356.3 | 1474.6 | -659.15 | 1318.3 | 16.99 | 11 | **0.10** |
| ***Fixed effect*** | *Odds ratio* | *95% CI* | *z* | *p* | | | |
| Int | 60.69 | 21.40 – 172.12 | 7.72 | **<0.001** | | | |
| Ips-Con | 0.64 | 0.30 – 1.39 | -1.12 | 0.263 | | | |
| Pr-Po | 0.62 | 0.29 – 1.34 | -1.21 | 0.225 | | | |
| Pul-Sha | 0.63 | 0.25 – 1.57 | -0.99 | 0.323 | | | |
| Cen-Sha | 0.97 | 0.35 – 2.72 | -0.05 | 0.959 | | | |
| Con-Ips:Pr-Po | 1.09 | 0.39 – 3.05 | 0.17 | 0.862 | | | |
| Con-Ips:Pul-Sha | 1.23 | 0.43 – 3.52 | 0.39 | 0.697 | | | |
| Con-Ips:Cen-Sha | 1.55 | 0.48 – 5.02 | 0.73 | 0.464 | | | |
| Pr-Po:Pul-Sha | 1.20 | 0.44 – 3.27 | 0.37 | 0.715 | | | |
| Pr-Po:Cen-Sha | 1.09 | 0.36 – 3.30 | 0.15 | 0.878 | | | |
| Con-Ips:Pr-Po:Pul-Sha | 2.29 | 0.54 – 9.72 | 1.12 | 0.261 | | | |

| | | | | | | |
|---|---|---|---|---|---|---|
| Con-Ips:Pr-Po:Cen-Sha | 0.51 | 0.11 – 2.33 | -0.87 | 0.382 | | |
| ***Random effect*** | *N (obs)* | *Factor* | *Variance* | *Std* | *Corr* | |
| Participant | 27 (3729) | Int | 4.20 | 3.31 | | |
| | | Pul-Sha | 1.00 | 1.06 | 0.44 | |
| | | Cen-Sha | 1.48 | 1.87 | 0.11 | 0.9 |
| Task order | | Int | 1.00 | 1.00 | | |

**Units in odds ratios. Abbreviations: Int, intercept; Ips, ipsilateral to the ultrasound target; Con, contralateral to the ultrasound target; Pr, pre-sonication; Po, post-sonication; Sha, sham sonication; Pul, pulvinar sonication; Cen, central thalamus sonication. Interaction terms include an ':' and represent the additional change in the included steps across levels of the variable.*

### Table 11: Correct responses on catch trials marginal means contrasts

| Contrast | Odds ratio | CI (lower) | CI (upper) | z | *p* | $p_{adj}$ |
|---|---|---|---|---|---|---|
| Pul-Sha \| Ips \| Po-Pr | 1.2 | 0.44 | 3.27 | 0.37 | 0.715 | 0.878 |
| Cen-Sha \| Ips \| Po-Pr | 1.09 | 0.36 | 3.3 | 0.15 | 0.878 | 0.878 |
| Cen-Pul \| Ips \| Po-Pr | 0.91 | 0.33 | 2.52 | -0.19 | 0.85 | 0.878 |
| Pul-Sha \| Con \| Po-Pr | 2.76 | 0.97 | 7.87 | 1.9 | 0.058 | 0.173 |
| Cen-Sha \| Con \| Po-Pr | 0.55 | 0.19 | 1.59 | -1.1 | 0.272 | 0.46 |
| Cen-Pul \| Con \| Po-Pr | 0.2 | 0.06 | 0.62 | -2.78 | **0.005** | **0.045** |
| Pul \| Con-Ips \| Po-Pr | 2.51 | 0.9 | 6.95 | 1.77 | 0.077 | 0.173 |
| Cen \| Con-Ips \| Po-Pr | 0.55 | 0.18 | 1.72 | -1.02 | 0.307 | 0.46 |
| Cen-Pul \| Con-Ips \| Po-Pr | 0.22 | 0.05 | 1.02 | -1.94 | 0.052 | 0.173 |

**Units in odds ratios. Abbreviations: Min, minute; Pr, pre-sonication; Po, post-sonication; S, sham; P, pulvinar; C, central thalamus.*

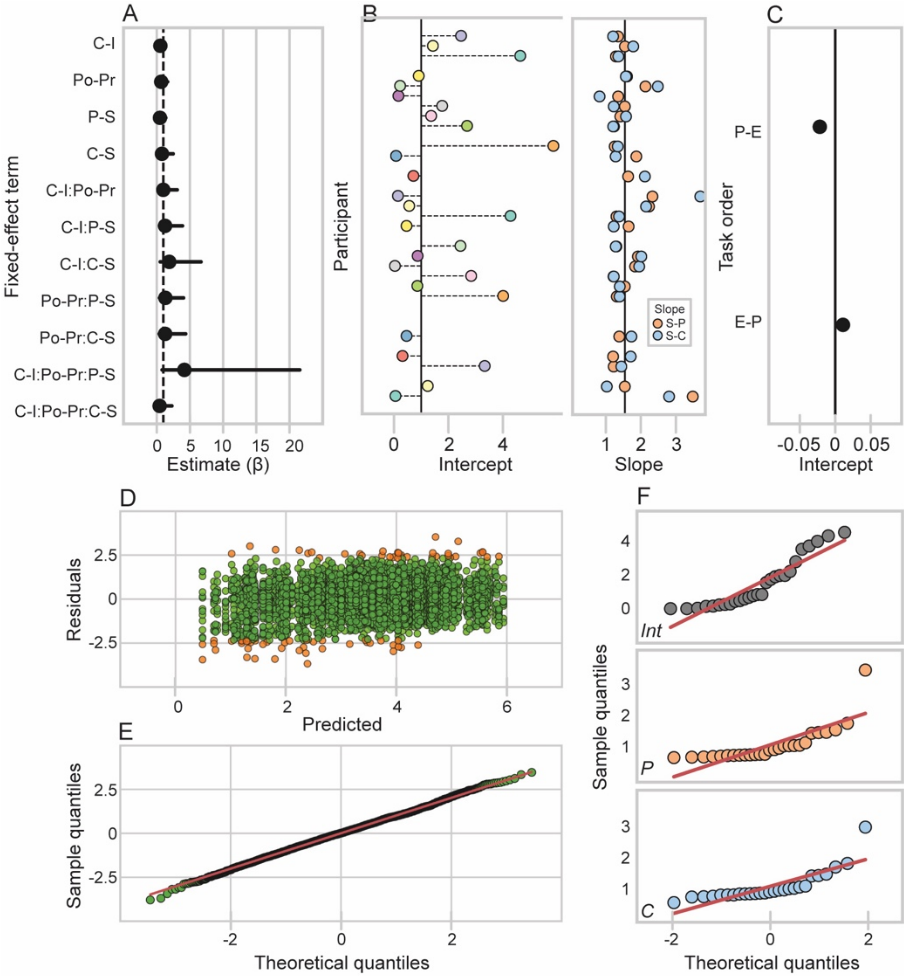


***Supplemental Figure 11:*** *Mixed-effects logistic regression for correct responses on catch trials ('correct rejections') during the Edgly-Driver Task (EDT). Fixed effects. Bars show 95% confidence intervals. (B & C): Random effects. (B): Random intercept for participant (left) with a random slope for ultrasound condition (right). The random intercept for each participant shows their baseline odds of responding correctly ('correct rejection') when block is at pre-sonication and ultrasound condition is at sham relative to the average baseline odds of responding correctly across participants estimated in the fixed effects, which is marked by the black vertical at 0. The random slope for ultrasound condition for each participant shows the difference in their baseline odds of responding correctly (when block is at pre-sonication) between the pulvinar and sham sonication conditions (orange) as well as between the central thalamus and sham sonication conditions (blue) relative to the average across participants estimated in the fixed effect, which is marked by the black vertical line at 0. (C): Random intercept for task order. Points show the average baseline odds of responding correctly (when block is at pre-sonication and ultrasound condition is at sham) for each task order group, specifically PVT then EDT (P-E) and EDT then PVT (E-P). (D): Predicted values versus residuals (simulated and normalizd using the DHARMa package in R). (E): QQ plot for theoretical quantiles versus simulated (and normalized) sample residual quantiles. (F): QQ plot showing theoretical quantiles versus sample quantiles for the random intercept for participant (top) as well as the random slopes for the steps in condition from sham to pulvinar sonication (middle) and from sham to central thalamus sonication (bottom).*

***Main trials.*** To examine whether sonication in any of the ultrasound conditions affected correct responses on main trials during the EDT ('hits'), and if this depended on cue validity and the visual field that the visual target appeared in, we fit a mixed-effect logistic regression testing the three-way interaction between the effects of ultrasound condition (sham, pulvinar, central thalamus sonication), block (before and after sonication), and cue-target location on accuracy: accuracy ~ condition × block × cue-target location + (1 + condition | participant) + (1 | task order). Cue-target location was a categorical variable with four levels: validly cued ipsilateral targets, validly cued contralateral targets, invalidly cued ipsilateral targets, and invalidly cued contralateral targets (see the Supplemental Material Fig. 9). One participant surpassed the Cook's distance threshold and underwent further evaluation (see Fig. 10D). This participant showed an erroneous change in correct responses across blocks during their sham session (see Fig. 10E). We excluded that session for that participant from the mixed-effect model. All other data were used in the model. Participants showed some variation in their baseline (pre-sonication) probability of responding correctly on main trials ('hits') between the ultrasound sessions, as indicated by the random slopes for condition for each participant (see Table 12 and Fig. 11B, right). There was a no meaningful variability in the probability of responding correctly on main trials between the task order groups (see Table 12 and Fig. 11C). All estimated marginal means contrasts performed are presented in Table 13. Follow-up estimated marginal means contrasts were used to make more specific comparisons in the models and adjusted for multiple comparisons using the False Discovery Rate (FDR).

**Table 12: Mixed-effect logistic regression: catch trial accuracy ~ spatial cue location × block × condition + (1 + condition | participant) + (1 | task order)**

**Model summary**

| | *Marg $R^2$* | *Cond $R^2$* |
|---|---|---|
| | 0.010 | 0.07 |

| ***Fit (ANOVA)*** | *AIC* | *BIC* | *Log likelihood* | *Deviance* | $\chi^2$ | *df* | *p* |
|---|---|---|---|---|---|---|---|
| Null/restricted | 29105 | 29172 | -14545 | 29089 | | | |
| Full | 28988 | 29246 | -14463 | 28926 | 163.2 | 23 | **<0.001** |

| ***Fixed effect*** | *Estimate* | *95% CI* | *t* | *p* |
|---|---|---|---|---|
| Int | 3.63 | 2.67 – 4.94 | 8.21 | **<0.001** |
| iIps-iCon | 0.83 | 0.59 – 1.17 | -1.08 | 0.281 |
| vCon-iCon | 0.91 | 0.69 – 1.20 | -0.67 | 0.506 |
| vIps-iCon | 1.35 | 1.02 – 1.78 | 2.09 | **0.037** |
| Pr-Po | 1.14 | 0.79 – 1.64 | 0.70 | 0.482 |
| Sha-Pul | 1.26 | 0.82 – 1.92 | 1.06 | 0.289 |
| Sha-Cen | 1.06 | 0.71 – 1.60 | 0.30 | 0.767 |
| iIps-iCon:Pr-Po | 1.09 | 0.67 – 1.76 | 0.35 | 0.727 |
| vCon-iCon:Pr-Po | 1.28 | 0.87 – 1.89 | 1.24 | 0.213 |
| vIps-iCon:Pr-Po | 1.21 | 0.81 – 1.79 | 0.94 | 0.349 |
| iIps-iCon:Sha-Pul | 0.90 | 0.55 – 1.47 | -0.44 | 0.661 |
| vCon-iCon:Sha-Pul | 0.94 | 0.63 – 1.41 | -0.30 | 0.766 |
| vIps-iCon:Sha-Pul | 0.78 | 0.52 – 1.17 | -1.19 | 0.235 |
| iIps-iCon:Sha-Cen | 1.03 | 0.64 – 1.67 | 0.13 | 0.900 |
| vCon-iCon:Sha-Cen | 1.18 | 0.80 – 1.74 | 0.85 | 0.397 |
| vIps-iCon:Sha-Cen | 1.00 | 0.68 – 1.48 | 0.00 | 0.996 |

| | | | | | | |
|---|---|---|---|---|---|---|
| Pr-Po:Sha-Pul | 0.78 | 0.46 – 1.32 | -0.92 | 0.358 | | |
| Pr-Po:Sha-Cen | 1.13 | 0.67 – 1.91 | 0.47 | 0.637 | | |
| iIps-iCon:Pr-Po:Sha-Pul | 1.26 | 0.63 – 2.53 | 0.66 | 0.512 | | |
| vCon-iCon:Pr-Po:Sha-Pul | 1.01 | 0.58 – 1.78 | 0.04 | 0.970 | | |
| vIps-iCon:Pr-Po:Sha-Pul | 1.13 | 0.64 – 1.99 | 0.41 | 0.683 | | |
| iIps-iCon:Pr-Po:Sha-Cen | 0.67 | 0.34 – 1.33 | -1.15 | 0.249 | | |
| vCon-iCon:Pr-Po:Sha-Cen | 0.61 | 0.35 – 1.07 | -1.73 | 0.083 | | |
| vIps-iCon:Pr-Po:Sha-Cen | 0.61 | 0.35 – 1.07 | -1.72 | 0.085 | | |
| ***Random effect*** | *N (obs)* | *Factor* | *Variance* | *Std* | *Corr* | |
| Participant | 27 (31649) | Int | 1.22 | 1.56 | | |
| | | Pul-Sha | 1.28 | 1.63 | -0.61 | |
| | | Cen-Sha | 1.27 | 1.63 | -0.46 | 0.76 |
| Task order | | Int | 1.00 | 1.00 | | |

** Units in log odds. 1 indicates equal probability of a correct response between groups compared. Abbreviations: Marg, marginal; Cond, conditional; iIps, invalidly cued ipsilateral target; iCon, invalidly cued contralateral target; vIps, validly cued ipsilateral target; vCon, validly cued contralateral target; Pr, pre-sonication; Po, post-sonication; Sha, sham sonication; Pul, pulvinar sonication; Cen, central thalamic sonication. Interaction terms represent the additional change in the outcome on the steps between the listed factors. Rows are italicized if they appeared in the main text.*

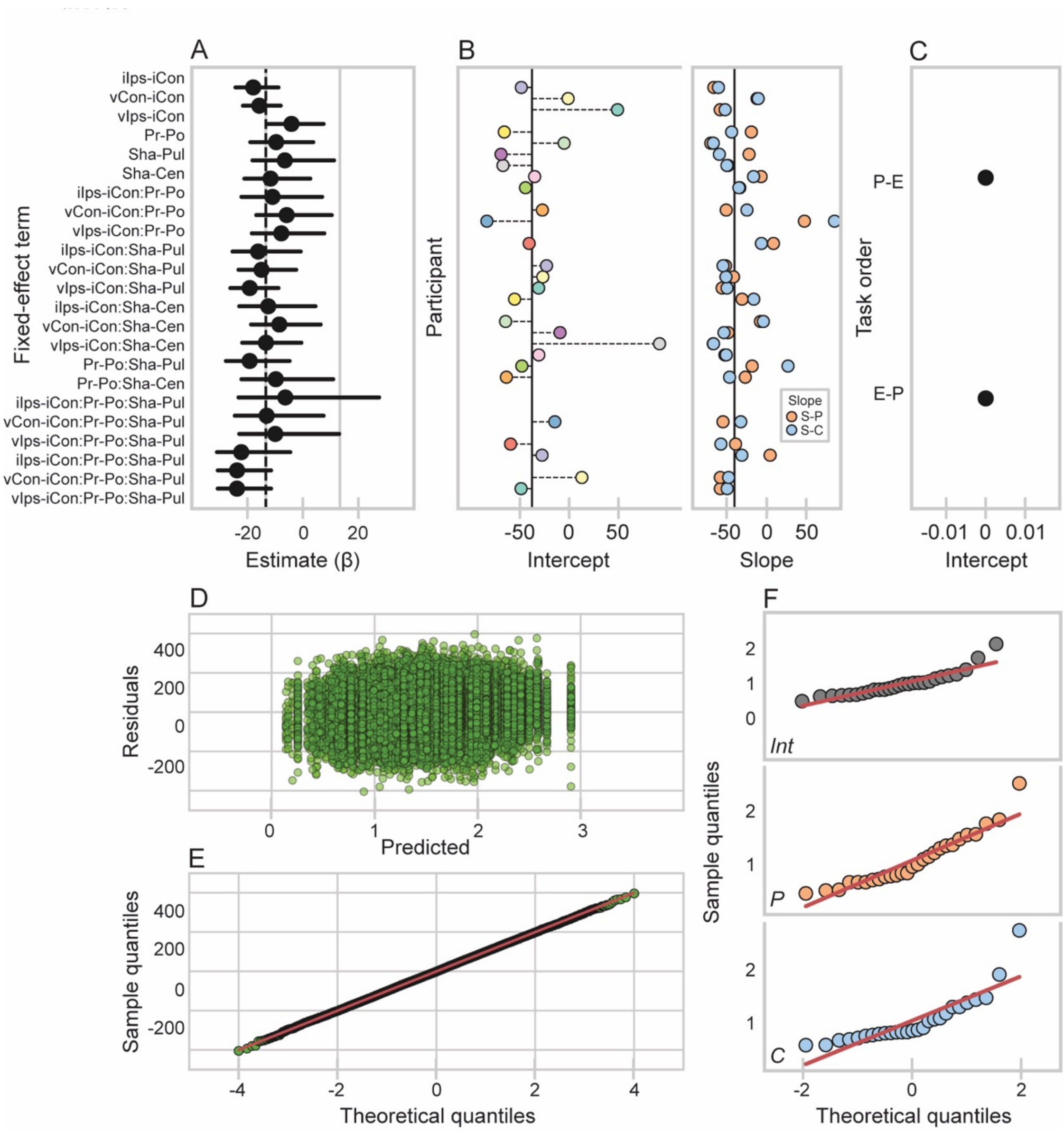


***Supplemental Figure 12:*** *Mixed-effects logistic regression for correct responses on main trials ('hits') during the Edgly-Driver Task (EDT). Fixed effects. Bars show 95% confidence intervals. (B & C): Random effects. (B): Random intercept for participant (left) with a random slope for ultrasound condition (right). The random intercept for each participant shows their baseline odds of responding correctly (when block is at pre-sonication and ultrasound condition is at sham) relative to the average baseline odds of responding correctly across participants estimated in the fixed effects (black vertical at 0). The random slope for ultrasound condition for each participant shows the difference in their baseline odds of responding correctly (when block is at pre-sonication) between the pulvinar and sham sonication conditions (orange) as well as between the central thalamus and sham sonication conditions (blue) relative to the average across participants estimated in the fixed effect (black vertical line at 0). (C): Random intercept for task order. Points show the average baseline odds of responding correctly (when block is at pre-sonication and ultrasound condition is at sham) for each task order group, PVT then EDT (P-E) and EDT then PVT (E-P). (D): Predicted values versus residuals (simulated and normalized using the DHARMa package in R). (. (E): QQ plot for theoretical quantiles versus simulated (and normalized) sample residual quantiles. (F): QQ plot showing theoretical quantiles versus sample quantiles for the*

*random intercept for participant (top) as well as the random slopes for the steps in condition from sham to pulvinar sonication (middle) and from sham to central thalamus sonication (bottom).*

**Table 15: Main trial accuracy estimated marginal means contrasts**

| Contrast | Estimate | CI (lower) | CI (upper) | z | p | $p_{adj}$ |
|---|---|---|---|---|---|---|
| Cen-Pul \| Po-Pr \| iCon | 1.45 | 0.83 | 2.53 | 1.31 | 0.19 | 0.433 |
| Pul-Sha \| Po-Pr \| iIps | 0.99 | 0.61 | 1.58 | -0.06 | 0.956 | 0.989 |
| Cen-Sha \| Po-Pr \| iIps | 0.76 | 0.48 | 1.2 | -1.18 | 0.24 | 0.433 |
| Cen-Pul \| Po-Pr \| iIps | 0.77 | 0.48 | 1.23 | -1.1 | 0.272 | 0.433 |
| Pul-Sha \| Po-Pr \| vCon | 0.79 | 0.64 | 0.98 | -2.19 | 0.029 | 0.225 |
| Cen-Sha \| Po-Pr \| vCon | 0.69 | 0.56 | 0.85 | -3.45 | 0.001 | 0.015 |
| Cen-Pul \| Po-Pr \| vCon | 0.88 | 0.71 | 1.08 | -1.26 | 0.209 | 0.433 |
| Pul-Sha \| Po-Pr \| vIps | 0.88 | 0.7 | 1.11 | -1.09 | 0.274 | 0.433 |
| Cen-Sha \| Po-Pr \| vIps | 0.69 | 0.55 | 0.87 | -3.21 | 0.001 | 0.015 |
| Cen-Pul \| Po-Pr \| vIps | 0.79 | 0.63 | 0.98 | -2.17 | 0.03 | 0.225 |
| Pul \| Po-Pr \| iIps - iCon | 1.38 | 0.82 | 2.32 | 1.2 | 0.23 | 0.433 |
| Pul \| Po-Pr \| vCon - iCon | 1.29 | 0.85 | 1.98 | 1.2 | 0.231 | 0.433 |
| Pul \| Po-Pr \| vIps - iCon | 1.36 | 0.89 | 2.08 | 1.42 | 0.156 | 0.433 |
| Cen \| Po-Pr \| iIps - iCon | 0.73 | 0.44 | 1.21 | -1.23 | 0.219 | 0.433 |
| Cen \| Po-Pr \| vCon - iCon | 0.78 | 0.52 | 1.18 | -1.16 | 0.246 | 0.433 |
| Cen \| Po-Pr \| vIps - iCon | 0.74 | 0.49 | 1.12 | -1.43 | 0.152 | 0.433 |
| Cen-Pul \| Po-Pr \| iIps - iCon | 0.53 | 0.26 | 1.09 | -1.72 | 0.086 | 0.36 |
| Cen-Pul \| Po-Pr \| vCon - iCon | 0.6 | 0.33 | 1.09 | -1.67 | 0.096 | 0.36 |
| Pul-Sha \| Po-Pr \| vCon - iIps | 0.8 | 0.48 | 1.34 | -0.84 | 0.4 | 0.6 |
| Cen-Sha \| Po-Pr \| vCon - iIps | 0.91 | 0.55 | 1.52 | -0.34 | 0.73 | 0.811 |
| Cen-Pul \| Po-Pr \| vCon - iIps | 1.14 | 0.68 | 1.91 | 0.51 | 0.613 | 0.8 |
| Pul-Sha \| Po-Pr \| vIps - iCon | 1.13 | 0.64 | 1.99 | 0.41 | 0.683 | 0.811 |
| Cen-Sha \| Po-Pr \| vIps - iCon | 0.61 | 0.35 | 1.07 | -1.72 | 0.085 | 0.36 |
| Cen-Pul \| Po-Pr \| vIps - iCon | 0.54 | 0.3 | 0.98 | -2.02 | 0.044 | 0.264 |
| Pul-Sha \| Po-Pr \| vIps - iIps | 0.89 | 0.53 | 1.51 | -0.43 | 0.67 | 0.811 |
| Cen-Sha \| Po-Pr \| vIps - iIps | 0.91 | 0.55 | 1.53 | -0.34 | 0.73 | 0.811 |
| Cen-Pul \| Po-Pr \| vIps - iIps | 1.02 | 0.61 | 1.72 | 0.09 | 0.93 | 0.989 |
| Pul-Sha \| Po-Pr \| vIps - vCon | 1.11 | 0.82 | 1.52 | 0.69 | 0.493 | 0.672 |
| Cen-Sha \| Po-Pr \| vIps - vCon | 1 | 0.74 | 1.36 | -0.01 | 0.993 | 0.993 |
| Cen-Pul \| Po-Pr \| vIps - vCon | 0.9 | 0.66 | 1.21 | -0.72 | 0.474 | 0.672 |

**Units in ms. Min (slope) represents the average change per minute into the task. Min (0) represents 0 minutes into the task considered task onset. Abbreviations: Min, minute; Pr, pre-sonication; Po, post-sonication; Sha, sham sonication; Pul, pulvinar sonication; Cen, central thalamic sonication.*

**Response time.** One linear mixed-effects model was used to test the three-way interaction between ultrasound condition (sham, pulvinar, central thalamus sonication), block (before and after sonication), and cue-target location on response times from main trials: response time ~ condition × block × cue-target location + (1 + condition | participant) + (1 | task order). Cue-target location was a categorical variable with four levels: validly cued ipsilateral visual targets, validly cued contralateral targets, invalidly cued ipsilateral targets, and invalidly cued contralateral targets (see Supplemental Fig 9). No participants surpassed the Cook's distance threshold (see Fig. 13C), so all data were used in the model. Participants showed some variation in their baseline (pre-sonication) response times on main trials during the EDT between the ultrasound sessions, as indicated by the random slopes for condition (see Table 14 and Fig. 13B, right). However, there was no meaningful variation in response times on main trials between the task order groups (see Table 14 and Fig. 13C), which suggests that the order in which participants completed the did could not explain the variation in response times during the EDT. All estimated marginal means contrasts performed are presented in Table 15. Follow-up estimated marginal means contrasts were used to make more specific comparisons in the models and adjusted for multiple comparisons using the False Discovery Rate (FDR).

**Table 14: Linear mixed-effect model: response time ~ cue-target location × block × condition + (1 + condition | participant) + (1 | task order)**

**Model summary**

| | *Marg $R^2$* | *Cond $R^2$* |
|---|---|---|
| | 0.03 | 0.19 |

| ***Fit (ANOVA)*** | *AIC* | *BIC* | *Log likelihood* | *Deviance* | *$\chi^2$* | *df* | *p* |
|---|---|---|---|---|---|---|---|
| Null/restricted | 279212 | 279285 | -139597 | 279194 | | | |
| Full | 278361 | 278622 | -139148 | 278297 | 897.1 | 23 | **<0.001** |

| ***Fixed effect*** | *Estimate* | *SE* | *95% CI* | *t* | *p* |
|---|---|---|---|---|---|
| Int | 359.61 | 5.82 | 348.20 – 371.03 | 61.76 | **<0.001** |
| iIps-iCon | 5.70 | 4.67 | -3.45 – 14.85 | 1.22 | 0.222 |
| vCon-iCon | -19.72 | 3.68 | -26.93 – -12.52 | -5.36 | **<0.001** |
| vIps-iCon | -13.60 | 3.67 | -20.78 – -6.42 | -3.71 | **<0.001** |
| Pr-Po | -12.26 | 4.84 | -21.75 – -2.77 | -2.53 | **0.011** |
| Sha-Pul | -4.81 | 6.67 | -17.88 – 8.25 | -0.72 | 0.470 |
| Sha-Cen | 0.78 | 6.15 | -11.28 – 12.83 | 0.13 | 0.900 |
| iIps-iCon:Pr-Po | -0.77 | 6.55 | -13.60 – 12.06 | -0.12 | 0.906 |
| vCon-iCon:Pr-Po | 0.16 | 5.18 | -9.98 – 10.31 | 0.03 | 0.975 |
| vIps-iCon:Pr-Po | -1.15 | 5.16 | -11.26 – 8.96 | -0.22 | 0.823 |
| iIps-iCon:Sha-Pul | 10.48 | 6.66 | -2.57 – 23.53 | 1.57 | 0.115 |
| vCon-iCon:Sha-Pul | 1.76 | 5.23 | -8.50 – 12.01 | 0.34 | 0.737 |
| vIps-iCon:Sha-Pul | 4.89 | 5.22 | -5.34 – 15.12 | 0.94 | 0.349 |
| iIps-iCon:Sha-Cen | -3.36 | 6.65 | -16.41 – 9.68 | -0.51 | 0.613 |
| vCon-iCon:Sha-Cen | -7.44 | 5.20 | -17.62 – 2.74 | -1.43 | 0.152 |
| vIps-iCon:Sha-Cen | -4.28 | 5.18 | -14.44 – 5.88 | -0.82 | 0.409 |
| Pr-Po:Sha-Pul | 15.30 | 6.96 | 1.66 – 28.95 | 2.20 | **0.028** |
| Pr-Po:Sha-Cen | 2.88 | 6.84 | -10.53 – 16.29 | 0.42 | 0.674 |
| iIps-iCon:Pr-Po:Sha-Pul | -10.85 | 9.36 | -29.19 – 7.48 | -1.16 | 0.246 |
| vCon-iCon:Pr-Po:Sha-Pul | -13.08 | 7.43 | -27.64 – 1.47 | -1.76 | 0.078 |
| vIps-iCon:Pr-Po:Sha-Pul | -11.47 | 7.40 | -25.97 – 3.04 | -1.55 | 0.121 |
| iIps-iCon:Pr-Po:Sha-Cen | 2.24 | 9.30 | -16.00 – 20.47 | 0.24 | 0.810 |
| vCon-iCon:Pr-Po:Sha-Cen | -0.89 | 7.31 | -15.21 – 13.44 | -0.12 | 0.903 |
| vIps-iCon:Pr-Po:Sha-Cen | -1.53 | 7.29 | -15.81 – 12.75 | -0.21 | 0.834 |

| ***Random effect*** | *N (obs)* | *Factor* | *Variance* | *Std* | *Corr* | |
|---|---|---|---|---|---|---|
| Participant | 27 (25639) | Int | 596.00 | 24.42 | | |
| | | Pul-Sha | 538.80 | 12.21 | -0.54 | |
| | | Cen-Sha | 382.20 | 19.55 | -0.29 | 0.44 |

| | | | |
|---|---|---|---|
| Task order | Int | 0.00 | 0.00 |

**Model fit with REML. Units in ms. Abbreviations: Marg, marginal; Cond, conditional; iIps, invalidly cued ipsilateral target; iCon, invalidly cued contralateral target; vIps, validly cued ipsilateral target; vCon, validly cued contralateral target; Pr, pre-sonication; Po, post-sonication; Sha, sham sonication; Pul, pulvinar sonication; Cen, central thalamic sonication. Interaction terms represent the additional change in the outcome on the steps between the listed factors.*

### Table 15: Response time estimated marginal means contrasts

| Contrast | Estimate | SE | CI (lower) | CI (upper) | z | p | $p_{adj}$ |
|---|---|---|---|---|---|---|---|
| Cen-Pul \| Po-Pr \| iCon | -12.43 | 6.96 | -26.06 | 1.21 | -1.79 | 0.074 | 0.553 |
| Pul-Sha \| Po-Pr \| iIps | 4.45 | 6.25 | -7.81 | 16.7 | 0.71 | 0.477 | 0.930 |
| Cen-Sha \| Po-Pr \| iIps | 5.12 | 6.31 | -7.25 | 17.48 | 0.81 | 0.417 | 0.930 |
| Cen-Pul \| Po-Pr \| iIps | 0.67 | 6.33 | -11.73 | 13.07 | 0.11 | 0.916 | 0.930 |
| Pul-Sha \| Po-Pr \| vCon | 2.22 | 2.61 | -2.89 | 7.33 | 0.85 | 0.395 | 0.930 |
| Cen-Sha \| Po-Pr \| vCon | 1.99 | 2.59 | -3.09 | 7.07 | 0.77 | 0.443 | 0.930 |
| Cen-Pul \| Po-Pr \| vCon | -0.23 | 2.57 | -5.27 | 4.81 | -0.09 | 0.930 | 0.930 |
| Pul-Sha \| Po-Pr \| vIps | 3.84 | 2.52 | -1.11 | 8.78 | 1.52 | 0.129 | 0.553 |
| Cen-Sha \| Po-Pr \| vIps | 1.35 | 2.51 | -3.58 | 6.28 | 0.54 | 0.591 | 0.930 |
| Cen-Pul \| Po-Pr \| vIps | -2.49 | 2.50 | -7.39 | 2.42 | -0.99 | 0.320 | 0.930 |
| Pul \| Po-Pr \| iIps - iCon | -11.62 | 6.68 | -24.73 | 1.48 | -1.74 | 0.082 | 0.553 |
| Pul \| Po-Pr \| vCon - iCon | -12.92 | 5.33 | -23.36 | -2.48 | -2.43 | 0.015 | 0.255 |
| Pul \| Po-Pr \| vIps - iCon | -12.62 | 5.31 | -23.02 | -2.21 | -2.38 | 0.017 | 0.255 |
| Cen \| Po-Pr \| iIps - iCon | 1.47 | 6.61 | -11.49 | 14.43 | 0.22 | 0.824 | 0.930 |
| Cen \| Po-Pr \| vCon - iCon | -0.72 | 5.16 | -10.84 | 9.39 | -0.14 | 0.889 | 0.930 |
| Cen \| Po-Pr \| vIps - iCon | -2.68 | 5.15 | -12.77 | 7.41 | -0.52 | 0.603 | 0.930 |
| Cen-Pul \| Po-Pr \| iIps - iCon | 13.09 | 9.40 | -5.33 | 31.52 | 1.39 | 0.164 | 0.597 |
| Cen-Pul \| Po-Pr \| vCon - iCon | 12.20 | 7.42 | -2.34 | 26.73 | 1.64 | 0.100 | 0.553 |
| Pul-Sha \| Po-Pr \| vCon - iIps | -2.23 | 6.77 | -15.5 | 11.04 | -0.33 | 0.742 | 0.930 |
| Cen-Sha \| Po-Pr \| vCon - iIps | -3.13 | 6.81 | -16.48 | 10.23 | -0.46 | 0.646 | 0.930 |
| Cen-Pul \| Po-Pr \| vCon - iIps | -0.90 | 6.83 | -14.28 | 12.49 | -0.13 | 0.896 | 0.930 |
| Cen-Sha \| Po-Pr \| vIps - iCon | -1.53 | 7.29 | -15.81 | 12.75 | -0.21 | 0.834 | 0.930 |
| Cen-Pul \| Po-Pr \| vIps - iCon | 9.94 | 7.39 | -4.55 | 24.43 | 1.34 | 0.179 | 0.597 |
| Pul-Sha \| Po-Pr \| vIps - iIps | -0.61 | 6.74 | -13.82 | 12.60 | -0.09 | 0.928 | 0.930 |
| Cen-Sha \| Po-Pr \| vIps - iIps | -3.77 | 6.79 | -17.07 | 9.53 | -0.55 | 0.579 | 0.930 |
| Cen-Pul \| Po-Pr \| vIps - iIps | -3.15 | 6.80 | -16.48 | 10.18 | -0.46 | 0.643 | 0.930 |
| Pul-Sha \| Po-Pr \| vIps - vCon | 1.62 | 3.61 | -5.47 | 8.70 | 0.45 | 0.654 | 0.930 |
| Cen-Sha \| Po-Pr \| vIps - vCon | -0.64 | 3.60 | -7.69 | 6.41 | -0.18 | 0.859 | 0.930 |
| Cen-Pul \| Po-Pr \| vIps - vCon | -2.26 | 3.59 | -9.29 | 4.77 | -0.63 | 0.529 | 0.930 |

**Units in ms. Abbreviations: Pr, pre-sonication; Po, post-sonication; Sha, sham sonication; Pul, pulvinar sonication; Cen, central thalamic sonication; iIps, invalidly cued ipsilateral target; iCon, invalidly cued contralateral target; vIps, validly cued ipsilateral target; vCon, validly cued contralateral target.*

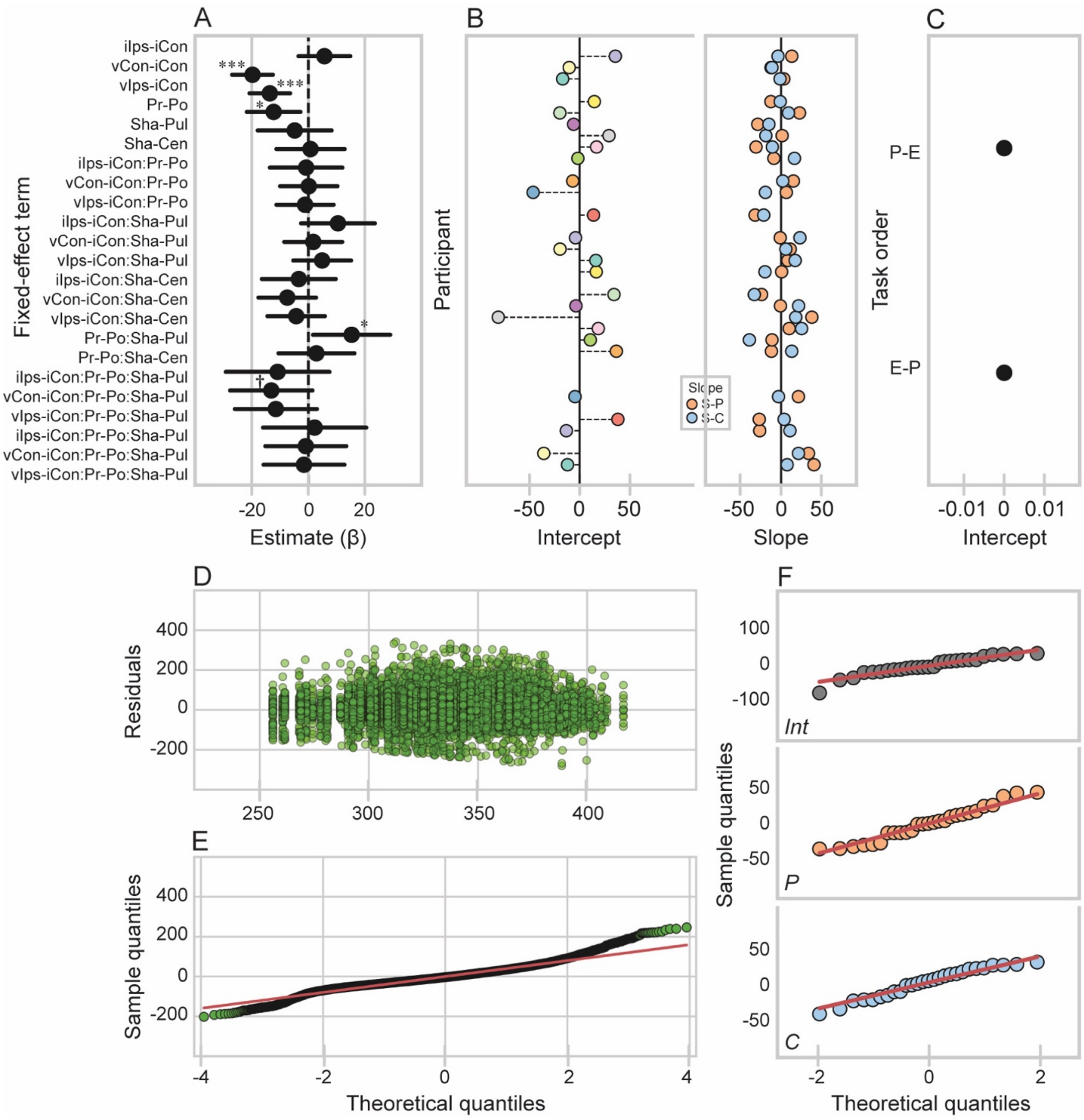


***Supplemental Figure 13:*** *Response time linear mixed-effect model for the Edgly-Driver Task (EDT). (A): Fixed effects. Bars show 95% confidence intervals. (B & C): Random effects. (B): Random intercept for participant (left) with a random slope for ultrasound condition (right). The random intercept for each participant shows their baseline response time (when block is at pre-sonication and ultrasound condition is at sham) relative to the average baseline response time across participants estimated in the fixed effects (black vertical at 0). The random slope for ultrasound condition for each participant shows the difference in their baseline response time (when block is at pre-sonication) between the pulvinar and sham sonication conditions (orange) as well as between the central thalamus and sham sonication conditions (blue) relative to the average across participants estimated in the fixed effect (black vertical line at 0). (C): Random intercept for task order. Points show the average baseline response time (when block is at pre-sonication and condition is at sham) for each task order group, PVT then EDT (P-E) and EDT then PVT (E-P). (D): Predicted values versus residuals. (E): QQ plot for theoretical quantiles versus residual (sample) quantiles. (F): QQ plot showing theoretical quantiles versus sample quantiles for the*

*random intercept for participant (top) as well as the random slopes for the steps in condition from sham to pulvinar sonication (middle) and from sham to central thalamus sonication (bottom).*

1. Tukey, J., *Exploratory Data Analysis Pearson.* London, United Kingdom.[Google Scholar], 1977.
2. Data, M.C., et al., *Noise versus outliers.* Secondary analysis of electronic health records, 2016: p. 163-183.
3. Benjamini, Y. and Y. Hochberg, *Controlling the False Discovery Rate: A Practical and Powerful Approach to Multiple Testing.* Journal of the Royal Statistical Society: Series B (Methodological), 1995. **57**(1): p. 289-300.
